# A Comprehensive Multi-Center Cross-platform Benchmarking Study of Single-cell RNA Sequencing Using Reference Samples

**DOI:** 10.1101/2020.03.27.010249

**Authors:** Wanqiu Chen, Yongmei Zhao, Xin Chen, Xiaojiang Xu, Zhaowei Yang, Yingtao Bi, Vicky Chen, Jing Li, Hannah Choi, Ben Ernest, Bao Tran, Monika Mehta, Malcolm Moos, Andrew Farmer, Alain Mir, Parimal Kumar, Urvashi Mehra, Jian-Liang Li, Wenming Xiao, Charles Wang

**Affiliations:** Center for Genomics, School of Medicine, Loma Linda University, 11021 Campus St., Loma Linda, CA 92350; CCR-SF Bioinformatics Group, Advanced Biomedical and Computational Sciences, Biomedical Informatics and Data Science Directorate, Frederick National Laboratory for Cancer Research, 8560 Progress Drive, Frederick, MD 21701; Department of Basic Sciences, School of Medicine, Loma Linda University, 11021 Campus St., Loma Linda, CA 92350; Integrative Bioinformatics Group, National Institute of Environment Health Sciences, NIH, Department of Health and Human Services, Research Triangle Park, NC 27709; Department of Allergy and Clinical Immunology, State Key Laboratory of Respiratory Disease, Guangzhou Institute of Respiratory Health, the First Affiliated Hospital of Guangzhou Medical University, Guangzhou, Guangdong, 510182, P.R. China; Abbvie Cambridge Research Center, 200 Sidney Street, Cambridge, MA 02139; Digicon Corporation, 7926 Jones Branch Drive, Suite 615, McLean, VA 22102; NCI CCR Sequencing Facility, Frederick National Laboratory for Cancer Research, Leidos Biomedical Research, Inc., 8560 Progress Drive, Frederick, MD 21701; Center for Biologics Evaluation and Research & Division of Cellular and Gene Therapies, U.S. Food and Drug Administration, 10903 New Hampshire Avenue, Silver Spring, MD 20993; Takara Bio USA, Inc., Mountain View, CA 94043; The Center for Devices and Radiological Health, U.S. Food and Drug Administration, Silver Spring, MA, 20993

## Abstract

Single-cell RNA sequencing (scRNA-seq) has become a very powerful technology for biomedical research and is becoming much more affordable as methods continue to evolve, but it is unknown how reproducible different platforms are using different bioinformatics pipelines, particularly the recently developed scRNA-seq batch correction algorithms. We carried out a comprehensive multi-center cross-platform comparison on different scRNA-seq platforms using standard reference samples. We compared six pre-processing pipelines, seven bioinformatics normalization procedures, and seven batch effect correction methods including CCA, MNN, Scanorama, BBKNN, Harmony, limma and ComBat to evaluate the performance and reproducibility of 20 scRNA-seq data sets derived from four different platforms and centers. We benchmarked scRNA-seq performance across different platforms and testing sites using global gene expression profiles as well as some cell-type specific marker genes. We showed that there were large batch effects; and the reproducibility of scRNA-seq across platforms was dictated both by the expression level of genes selected and the batch correction methods used. We found that CCA, MNN, and BBKNN all corrected the batch variations fairly well for the scRNA-seq data derived from biologically similar samples across platforms/sites. However, for the scRNA-seq data derived from or consisting of biologically distinct samples, limma and ComBat failed to correct batch effects, whereas CCA over-corrected the batch effect and misclassified the cell types and samples. In contrast, MNN, Harmony and BBKNN separated biologically different samples/cell types into correspondingly distinct dimensional subspaces; however, consistent with this algorithm’s logic, MNN required that the samples evaluated each contain a shared portion of highly similar cells. In summary, we found a great cross-platform consistency in separating two distinct samples when an appropriate batch correction method was used. We hope this large cross-platform/site scRNA-seq data set will provide a valuable resource, and that our findings will offer useful advice for the single-cell sequencing community.

## Introduction

Rapidly developing single-cell RNA sequencing (scRNA-seq) technologies allow interrogation of the transcriptome in unprecedented detail^1–3, 4^, but questions arise as to how accurate and reproducible different platforms are; and benchmarking and validation studies using standard reference samples have not appeared. Ziegenhain et al. reported a study comparing six different scRNA-seq library construction protocols and methods, but did not investigate the effects of bioinformatics factors such as different normalization and batch effect correction methods, not even mentioning that they only studied 583 mouse embryonic stem cells^5^. Because many of the factors influencing the results of single cell analyses-including stochastic events occurring during cell culture or cell isolation, single cell capture, library construction, and sequencing etc.^6^, remain to be identified, batch effects are a major, but underappreciated issue in scRNA-seq when comparisons need to be made between single-cell RNA-seq analyses within one laboratory or between laboratories^6, 7^. Batch effects may be derived from both technical and biological resources. The suitability and performance of limma and ComBat, the two batch correction algorithms originally developed for bulk cell RNA-seq data, for scRNA-seq data analyses have been questioned^7^. In 2018, four novel batch-effect-correction algorithms were reported for scRNA-seq data: Canonical Correlation Analysis (CCA)^7^, Mutual Nearest Neighbors (MNN)^6^, Scanorama^8, 9^, and Batch-Balanced k-Nearest Neighbors (BBKNN)^10, 11^ worked well for scRNA-seq batch effect corrections on scRNA-seq data under certain conditions^8, 10^. Tian et al. took an integrated computational analysis approach, evaluating three batch correction methods using four scRNA-seq datasets derived from two batches of mixtures of lung cancer lines from a single laboratory^12^. However, no systemic cross-platform comparison or performance evaluation of these newly developed batch correction algorithms using standard reference samples from which scRNA-seq data were generated across different centers has been reported.

Under the umbrella of the 2^nd^ phase of the FDA Sequencing Quality Control (SEQC-2) Consortium, we exploited a comprehensive study design including four scRNA-seq platforms: 10X Genomics Chromium, Fluidigm C1, Fluidigm C1 HT, and WaferGen, across five testing sites with seven different scRNA-seq data sets, using two well-characterized (see the companion manuscripts submitted to *Nature Biotechnology*), but biologically distinct reference cell lines, a human breast cancer cell line with a matched “normal” control cell line derived from B lymphocytes from the same patient^13^. Our goals were: 1) to evaluate the reproducibility of scRNA-seq across different platforms and sites using different bioinformatics pipelines and batch correction algorithms; 2) to establish benchmarking and quality control metrics for scRNA-seq; 3) to interrogate and benchmark certain cell type specific gene markers, in terms of the consistency across different scRNA-seq technologies and sites.

We also evaluated the effects of sequencing method and depth on the reproducibility of scRNA-seq and compared six different scRNA-seq data preprocessing pipelines, seven normalization methods, and seven different batch correction algorithms using our unique 20 scRNA-seq data sets derived from both mixed and non-mixed cell lines. Our analyses indicated that batch effects existed and the reproducibility of scRNA-seq across platforms and sites was dictated by both genes selected and bioinformatics pipelines, i.e. batch correction algorithms used. We found the performance of the newly developed CCA^7^, MNN^6^, Harmony^16, 17^, and BBKNN^10, 11^ batch correction algorithms depended on the nature of the biological samples or composition of cell types, consisting of either biologically similar/identical or distinct samples or cells. In summary, we found good cross-platform consistency in separating two distinct samples as long as an appropriate batch correction method was used.

## Results

### 1. Study design, overall data generated, and data QC assessments

**Figure 1a** shows our overall study design. It included four scRNA-seq platforms: 10X Genomics Chromium, Fluidigm C1, Fluidigm C1 HT, and WaferGen, across four testing sites (10X_LLU, 10X_NCI, C1_FDA_HT, C1_LLU, and WaferGen), using two well-characterized reference cell lines, a human breast cancer cell line (sample A) and a matched control normal B lymphocyte cell line (sample B) derived from the same patient^13^. Overall, we generated seven different scRNA-seq data sets including 4 different 3’-transcript scRNA-seq datasets (10X_LLU, 10X_NCI, 10X_NCI_M (modified shorter sequencing protocol), C1_FDA) and three different full-length transcript scRNA-seq datasets (C1_LLU, WaferGen_PE, and WaferGen_SE) (**Fig. 1a and Table 1**). For the 10X single-cell platform, we compared the standard sequencing protocol (26×98 bp) to the modified sequencing (26×56 bp) using the same scRNA-seq libraries. For the WaferGen platform, we also compared paired-end (75×2 bp) to single-end but much deeper sequencing (150 bp). For the scRNA-seq data, we applied 3 different pre-processing pipelines for the 3’-transcript scRNA-seq and three different pre-processing pipelines for the full-length scRNA-seq (**Suppl. Table 1 & 2**). We also evaluated seven different normalization methods and six different batch effect correction algorithms (**Fig. 1a**).

**Figure 1.**
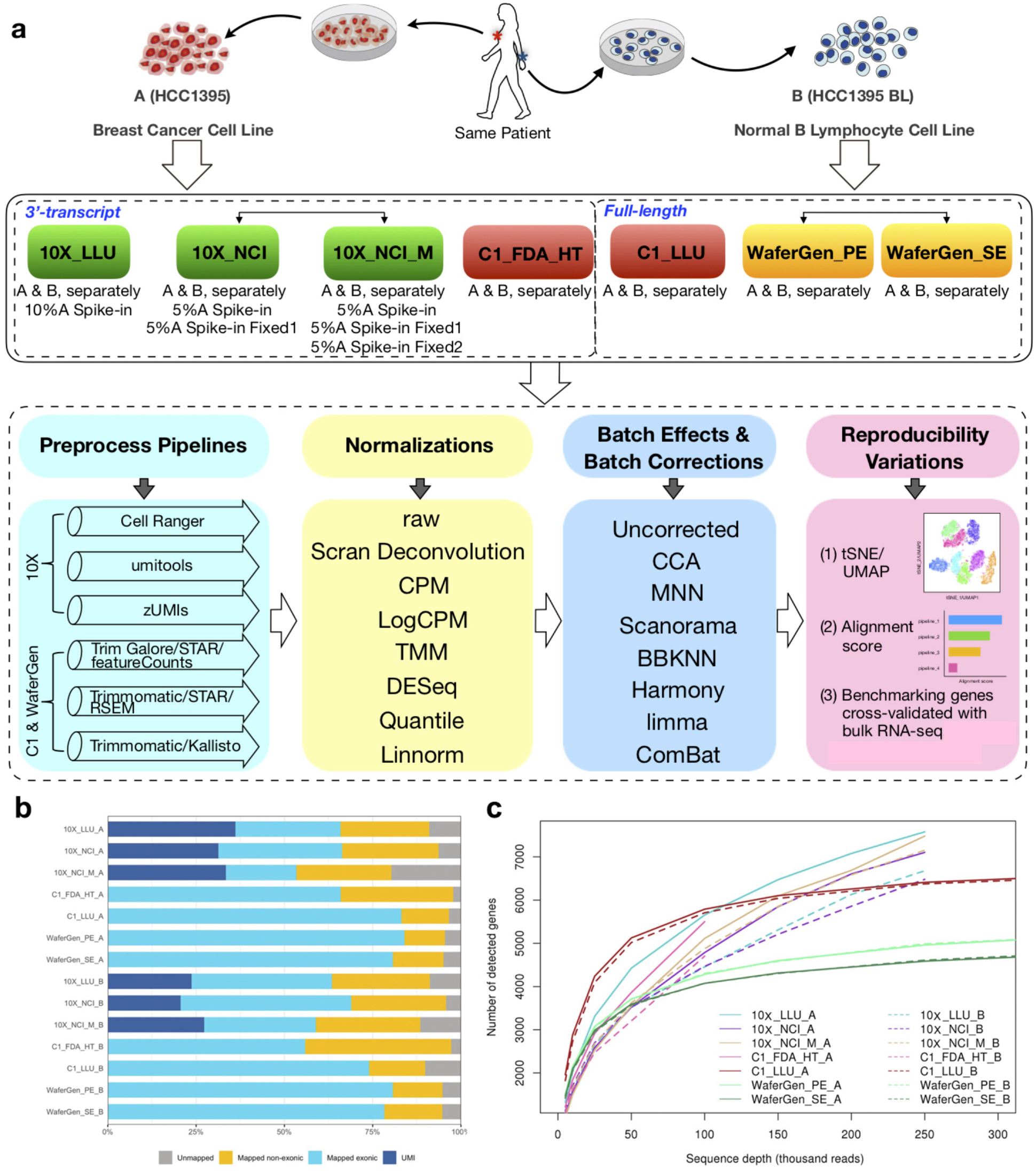
Overall study design and scRNA-seq mapping and numbers of detected genes across platforms. **(a)**. Schematic overview of the study design. Two well-characterized reference cell lines (sample A of a breast cancer cell line & sample B of a matched control normal B lymphocyte cell line) were used to generate scRNA-seq data across four platforms (10X Genomics, Fluidigm C1, Fluidigm C1 HT, and WaferGen), five testing sites (10X_LLU, 10X_NCI, C1_FDA_HT, C1_LLU, and WaferGen) using standard manufactures’ protocols. At 10X_LLU and 10X_NCI sites, mixed singe-cell capture and library constructions were also prepared with either 10% or 5% cancer cells spiked into B lymphocytes. At the NCI site, single-cell capture and library construction was also performed in fixed and mixed cells (5% cancer cell spiked into B lymphocytes). One set of 10X scRNA libraries from NCI was also sequenced using a shorter modified sequencing method. Bulk cell level RNA-seq data were also obtained from these cell lines, each in triplicate. All scRNA-seq data were subject to 3 different pre-processing pipelines for either 10X or C1/WaferGen technologies, respectively. We evaluated seven normalization methods such as Scran Deconvolution, CPM, LogCPM, TMM, DESeq, Quantile, and linnorm and seven batch effect correction algorithms including CCA, MNN, Scanorama, BBKNN, Harmony, limma, and ComBat. The cross-platform and cross-center performances were evaluated further by t-SNE, UMAP, modified alignment score, and both dot and feature plotting on certain selected marker genes. **Abbreviations and notations for** Fig. 1a**: 10X_LLU**, single cells were captured using 10X Genomics Chromium controller and scRNA-seq were sequenced at LLU Center for Genomics using the standard 10X Genomics protocol (26×98 bp); **10X_NCI_M**, 10X Genomics scRNA-seq libraries were prepared and sequenced at NCI sequencing facility using a modified 10X sequencing protocol (26×56 bp); **10X_NCI**, the same 10X Genomics scRNA-seq libraries were prepared at the NCI sequencing facility but sequenced at LLU using the standard 10X sequencing protocol (26×98 bp); **C1_FDA_HT**, single cells were captured using Fluidigm C1 HT IFC and the scRNA-seq libraries were sequenced at the FDA/CBER sequencing facility (75×2 bp); **C1_LLU**, single cells were captured using Fluidigm C1 IFC chip and the scRNA-seq libraries were sequenced at the LLU Center for Genomics (150×2 bp, ∼4-4.77M reads/cell); **WaterGen_PE**, single cells were captured using the ICELL8 chip (Takara Bio) and scRNA-seq libraries were sequenced at paired ends (75×2 bp) at Takara Bio; **WaterGen_SE**, the same scRNA-seq libraries generated at Takara Bio were sequenced at the LLU Center for Genomics (150×1 bp, ∼1M reads/cell). See **Table 1** for detail on the numbers of single cells captured and sequencing read depths in each platform and each site. **(b).** For both the breast cancer cell line (A) and normal B lymphocyte cell line (B) across 7 data sets, percentage of reads mapped to the exonic region (blue), non-exonic region (orange), or not mapped to the human genome (gray). For UMI methods (10X genomics platform), dark blue indicates the exonic reads with UMIs. **(c).** Median number of genes detected per cell at different sequencing read depth. Solid line represents the breast cancer cell line (A). Dashed line represents the normal B lymphocyte cell line (B).

**Table 1** summarizes the overall cell numbers and sequencing reads of single cells captured across four different sites, which provided seven different scRNA-seq data sets. A total of **25,265** single cells with whole transcriptomic scRNA-seq data were captured (**Table 1 & Suppl. Fig. 1**). Across all the platforms and data sets, over 94.0% of the reads were mapped to the exonic and non-exonic regions except for sample A of 10X_NCI_M (modified shorter sequencing), which had a mapping rate of 80.3% (sample A) and 88.5% (sample B) (**Fig. 1b**). However, there were variations in the mapping rates to exonic regions across platforms and sites, with WaferGen and Fluidigm C1 full-length transcript being higher (19.6%) than 3’-transcript scRNA data in tumor cells (A) (C1_LLU_A, 83.1%; WaferGen_PE_A, 84.0%; WaferGen_SE_A, 80.7%). The UMI (unique molecular identifier) data generated by the 10x platform showed that 34.8% of the exonic reads were derived from non-PCR amplified transcripts in tumor cells (sample A), and 26.4% of the exonic reads were derived from non-PCR amplified transcripts in normal B cells (sample B). We also noticed that the exonic mapping rates were slightly lower for the 10X genomics technologies when using a modified shorter modified sequencing protocol (26×56 bp vs. 26×98 bp). Nevertheless, many overlapping genes were detected (96.6%-97.3%) with a high correlation (R=0.997-0.998) between the normal and modified sequencing protocol for the 10X genomics scRNA-seq (**Suppl. Fig. 2**).

To investigate the effect of sequence depth on the number of genes detected across all platforms and scRNA-seq data sets, we down-sampled reads to varying depths on the data derived from all the platforms to assess the number of detected genes as well as the saturation level at the same sequence depth (**Fig. 1c**). We observed that for both tumor cells (A) and normal B-lymphocytes (B), the full-length cDNA transcript-based technologies (C1_LLU and WaferGen) displayed sequencing saturation at much lower sequence depth (∼50k) compared with 3’ scRNA-seq technologies, for which numbers of detected genes increased continuously with sequencing depth up to 250k reads (**Fig. 1c**).

For benchmarking scRNA-seq data, we determined and identified a large number of differentially expressed genes (DEGs) between the two cell lines at the population level (**Suppl. Data 1**, using fold-change (≥ 2) plus P-value (≤ 0.01, FDR= 0.05). **Supplemental Figure 3** shows the overall QC mapping of the population cell RNA-seq data for two cell lines (**Suppl. Fig. 3).**

### 2. Effects of data pre-processing

For the UMI based scRNA-seq data, we used three pipelines for preprocessing data: Cell Ranger (10X Genomics)^18^, UMI-tools^19^, and zUMIs^20^ to assess the consistency of the number of barcoded cells captured and the number of genes detected per cell (**Fig. 2a & 2b, Suppl. Table 1**). For the non-UMI based scRNA-seq data, we used three pre-processing pipelines: FeatureCounts^21^, Kallisto^22^, and RSEM^23^ to assess the consistency (**Fig. 2d & Suppl. Table 2**), which included trimming processes (cutadapt or trimmomatic), alignment (STAR and Kallisto), and gene counting (FeatureCounts, Kallisto, and RSEM). For simplicity, we used FeatureCounts, Kallisto, and RSEM to refer to the 3 non-UMI-based pipelines in this paper. We observed that, for the UMI-based scRNA-seq data, although both the number of cells and number of expressed genes per cell derived from each pre-processing pipeline were very similar, there were variations across three pipelines in all UMI based scRNA-seq data sets (**Fig. 2a & 2b**). Cell Ranger was the most conservative method for barcode cells selection. zUMIs showed the highest number of genes detected per cell. In addition, the gene expression level and the consensus genes per cell were highly correlated between any 2 pipelines for UMI-based pre-processing pipelines; Umi-tools and zUMIs showed the highest concordance.

**Figure 2.**
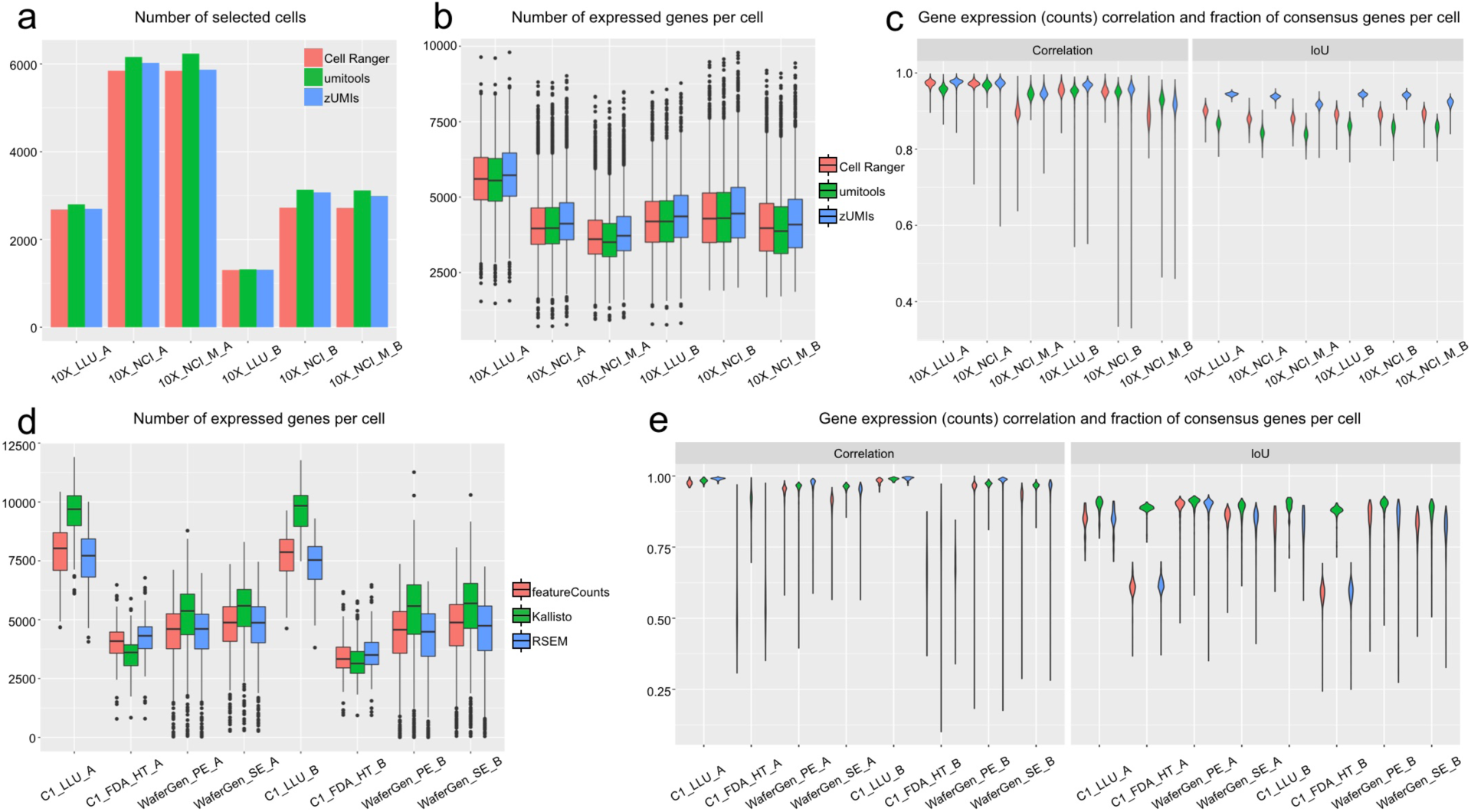
Effects of pre-processing pipelines on the number of genes detected with UMI-(a-c) and non-UMI-based (d-e) scRNA-seq platforms/data. **(a-c)** Effect of three pre-processing pipelines (Cell Ranger, UMI-Tools, zUMIs) on the UMI-based (10X) technology. Libraries from the two cell lines were constructed and sequenced by two sites (LLU and NCI). **(d-e)** Effect of three pre-processing pipelines (FeatureCounts, Kallisto, RSEM) on the non-UMI based technologies (C1 full transcript, C1 HT, and WaferGen full transcript). Libraries from the two cell lines were constructed and sequenced by three sites (C1_LLU, C1_FDA_HT, and WaferGen). **(a)** Barplot of the number of cells captured with UMI-based technology. **(b)** and **(d)** Boxplot of the number of genes detected per cell in UMI-based and non-UMI based technologies. **(c)** and **(e)** Violin plot of the gene expression correlation and consensus genes [represented by IoU (Intersection over Union)] per cell between any two pipelines in UMI-based and non-UMI based technologies.

For non-UMI based scRNA-seq data, much larger variation was observed in the number of genes detected across three pre-processing pipelines in all scRNA-seq data (**Fig. 2d**). Interestingly, we found that Kallisto identified a significantly higher number of genes per cell in the full-length transcript scRNA-seq, C1_LLU and WaferGen, whereas it detected of the fewest genes per cell in the C1-FDA_HT (3’ counting) dataset (**Fig. 2d**). In addition, the consensus genes per cell from the Kallisto pipeline varied significantly compared with the gene list from the other two pipelines for the C1-HT 3’ method, suggesting that the performance of alignment based (STAR) and alignment-free tools (Kallisto) might be inconsistent when preprocessing scRNA-seq data from 3’-based technologies. Overall, we found that the gene expression (counts) and the fraction of consensus genes per cell were highly variable across three pre-processing pipelines, both for the UMI- and non-UMI based scRNA-seq data (**Fig. 2c & 2e**). To simplify our comparison, we used the Cell Ranger for UMI-based and the FeaturesCounts for non-UMI-based pre-processing pipelines derived gene counts for all of our subsequent analyses.

### 3. Effects of normalizations

One special characteristic of scRNA-seq is sparsity of the data, which characteristically include high proportions of zero read counts^24^. The zero inflation can occur due to both biological (e.g., bi-stable gene regulation) and technical reasons (e.g., ‘drop out’ due to Poisson sampling limitations or limited efficiency of reverse transcription), which make the normalization of scRNA-seq data very challenging. Global scaling normalization methods developed for bulk RNA-seq data have been used for scRNA-seq data; these include CPM (Counts per Million), UQ (upper quantile), TMM (trimmed mean of M-value), and DESeq etc^25^. Regression based methods have also been proposed to remove the known nuisance factors in scRNA-seq data. It has been popular to regress out cell-cell variation in gene expression driven by cell alignment rate, the number of detected molecules, and mitochondrial gene expression after CPM normalization. There are also methods that are specifically tailored to scRNA-seq data sets, such as scran, SCnorm^26^, and Linnorm^27^ etc.

We used an algorithm, Silhouette width, to evaluate the performance of 7 different normalization methods: Scran deconvolution^28^, CPM, LogCPM, TMM, DESeq, quantile, and Linnorm (**Fig. 3a-g & 3h-n**). The Silhouette width is based on how well the two samples from the same cell are grouped with each other (**see Methods**). We observed good consistency in silhouette scores for all these methods both breast cancer cells and normal B-lymphocyte cells except for TMM and quantile, which failed to normalize the samples as they both had scores similar to the un-normalized raw data (**Fig. 3a-g & 3h-n**). A similar observation was confirmed using the 10X_LLU sample B dataset with different sequencing read depths (subsampled datasets, data not shown).

**Figure 3.**
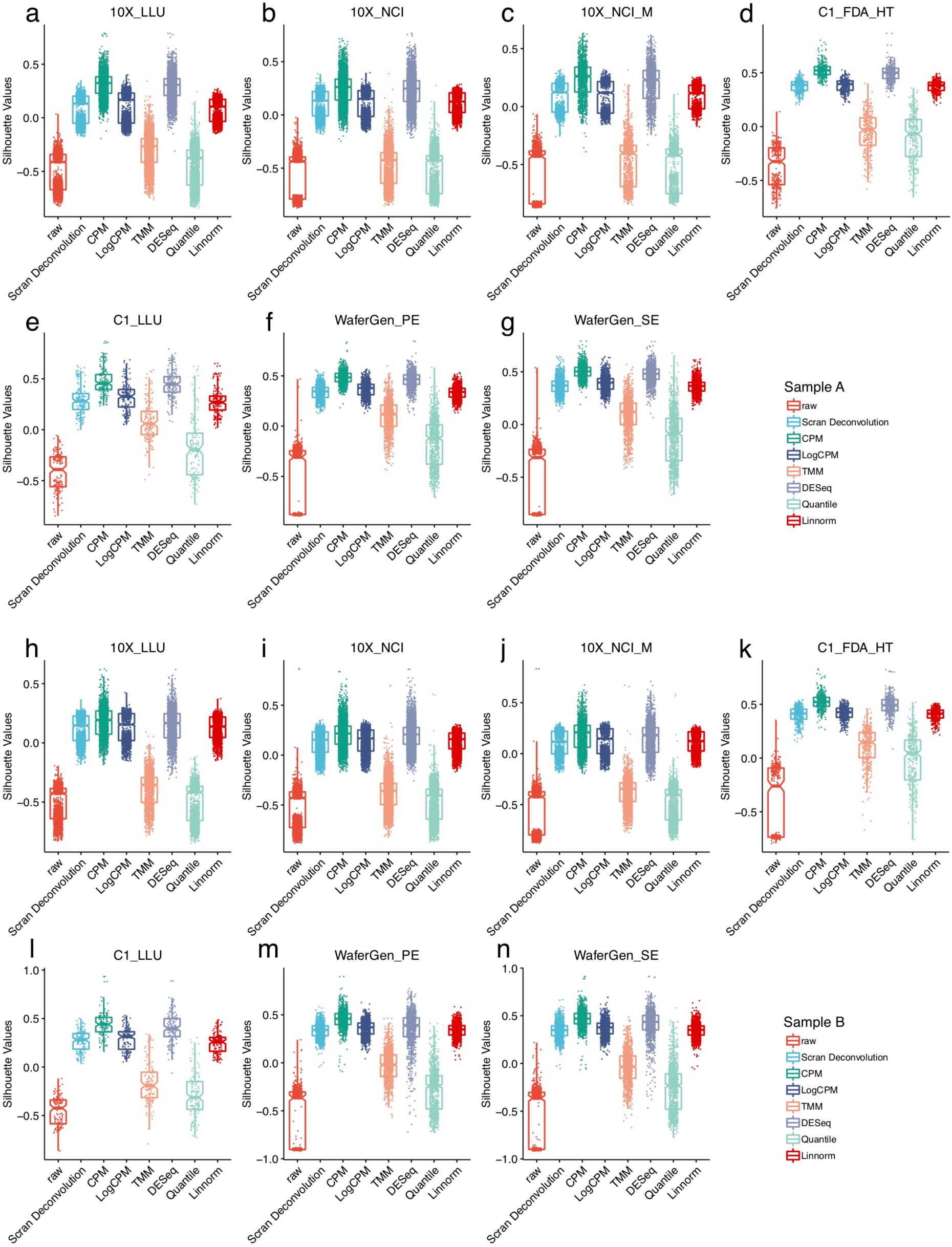
Silhouette scores of different normalizations in different scRNA-seq datasets. Evaluation of seven normalization methods, Scran Deconvolution, CPM, LogCPM, TMM, DESeq, Quantile and Linnorm with the silhouette width across different platforms and data sets. For each dataset, reads of each cell were downsampled to two different read depths (10K and 100K per cell) before calculating the silhouette width values. lgCPM is good enough for accurate clustering analysis. Two normalization methods developed for bulk RNA-seq had the lowest scores (TMM and Quantile). DESeq has similar performance as LogCPM **(a-g)** Boxplot of silhouette values stratified by seven normalization methods across seven datasets, including 10X_LLU (a), 10X_NCI (b), 10X_NCI_M (c), C1_FDA_HT (d), C1_LLU (e), Wafergen_PE (f) and WaferGen_SE (g) in breast cancer cells (sample A). **(h-n)** Boxplot of silhouette values stratified by seven normalization methods across seven datasets, including 10X_LLU (h), 10X_NCI (i), 10X_NCI_M (j), C1_FDA_HT (k), C1_LLU (l), Wafergen_PE (m) and WaferGen_SE (n) in normal B lymphocyte cells (sample B).

Distinguishing between unwanted variations and biologically significant changes can be difficult. A common step in preprocessing single-cell RNA-seq data is to regress out the so-called unwanted variations. However, we found that this did not improve the downstream clustering analysis (**Suppl. Fig. 4a-g & Suppl. Fig. 5a-g**). We also evaluated the consistency of Silhouette scores across different scRNA-seq platforms and data sets using Scan deconvolution, CPM, LogCPM, DESeq, and Linnorm, and we found two different patterns of performance: the 10X scRNA-seq data (across two sites and three different data sets) gave consistently lower scores than the C1 (both full-length and 3’ across two sites) and WaferGen (both SE and PE sequencing) (**Suppl. Fig. 6a-e & Suppl. Fig 7a-e**). It should be noted that fewer cells were used for the C1 (66 up to 200 cells) and WaferGen (∼600 cells) than for the 10X. Since log transformation has a high impact on downstream feature selection and clustering analysis and our analysis showed it performed fairly well, to simplify our comparison, we mainly used logCPM normalization in our subsequent batch effect and benchmarking evaluations except for the specific normalization methods embedded in some pipelines.

### 4. scRNA-seq data batch effects and batch correction

As noted earlier, batch effects can result from both technical and biological variations^24, 29^. Most existing normalization methods were developed for bulk RNA-seq^30^, so it was not surprising that normalization alone did not remove batch effects apparent in our data. For example, neither regressing mitochondrial genes nor normalizing UMI removed these effects (**Suppl. Fig. 8a/b**). We performed in-depth benchmarking evaluations using six batch effect correction algorithms: CCA^7^, MNN^6^, Scanorama^8, 9^, BBKNN^10, 11^, limma^31^, and ComBat^32^.

First, taking the gene counts based on the preprocessing pipelines selected above with the logCPM normalization using all scRNA-seq data across all sites and platforms, including spiking in samples (20 datasets), we applied these batch correction algorithms plus Harmony^16, 17^ to determine which one can separate the two different cell line samples correctly. Interestingly, we found that CCA overcorrected, and Scanorama, limma, and ComBat failed to separate cancer cells from B cells, whereas both MNN, Harmony and BBKNN worked well in separating cancer cells from B cells (**Suppl. Fig. 9a-h and Suppl. Data 2**).

Second, we evaluated these methods with scRNA-seq data derived from samples containing biologically similar cell types, i.e., sample A *or* sample B, analyzed separately (**Fig. 4a & b**), as well as samples where either 5% or 10% of cancer cells were spiked into the sample B cells, i.e., 10X_10%A_spikein_LLU, 10X_5%A_spikein_NCI, 10x_5%A_spikein_F1_NCI, 10X_5%A_spikein_F2_NCI, analyzed with the 10X Genomics platform across two sites (**Fig. 1a and Fig. 4c**). We performed batch-correction using each of the methods listed above on the top 1000 highly variable genes (HVG) on the data from all sites and platforms (**Fig. 4a-c**). We calculated a modified alignment score (**see Methods**) after each batch correction^7^. Without batch correction, t-SNE plotting showed that the cells from different batches or platforms/sites were clustered separately, not evenly mixed, indicating large variation and/or strong batch effects existed (**Fig. 4a-c, left panels**).

**Figure 4.**
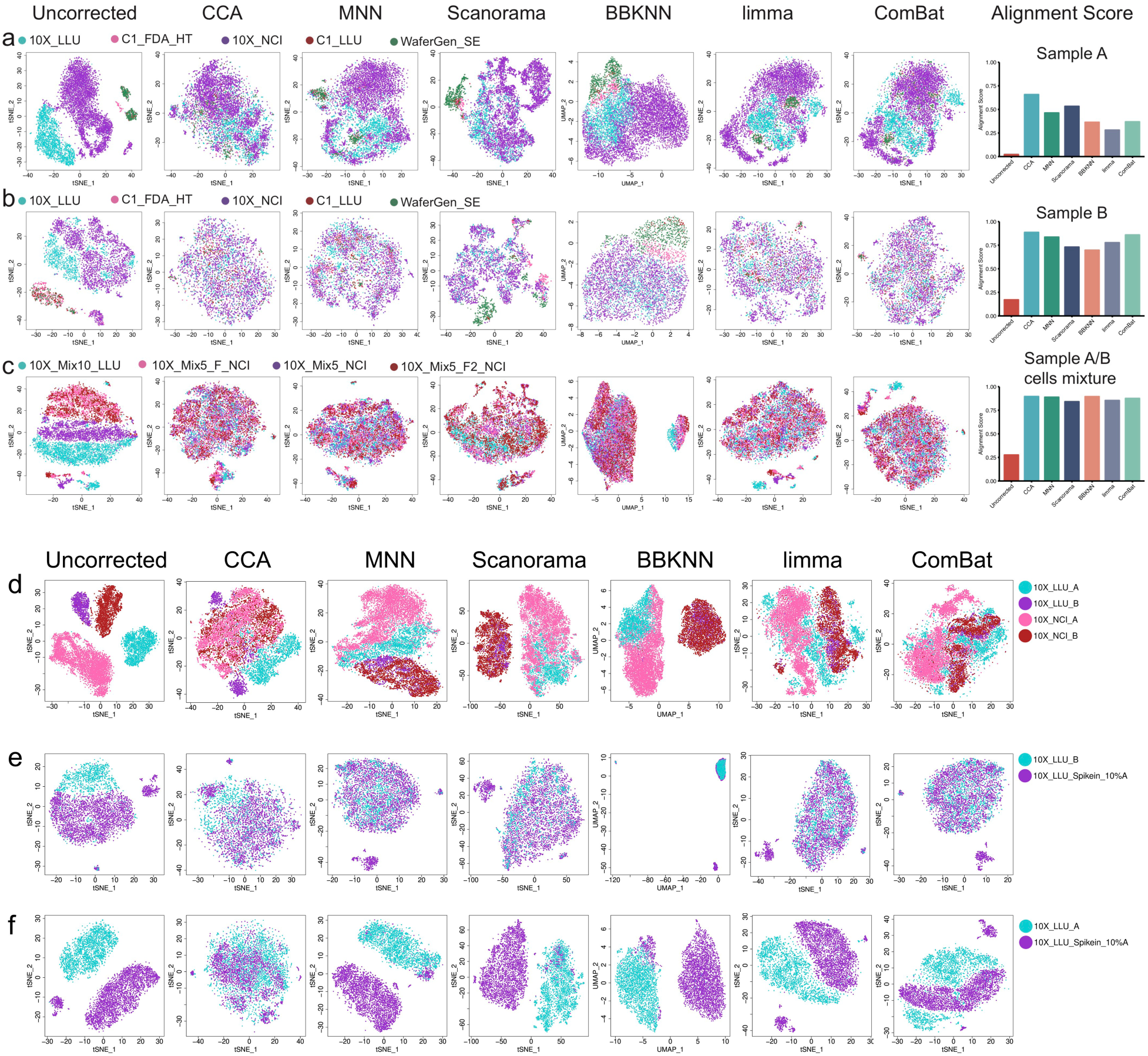
Evaluation of batch effect correction methods in different sample scenarios. **(a)-(b)** Batch effect corrections were performed using scRNA-seq data from five different sites and/or platforms in biologically similar cells, either breast cancer cells **(a)** or B lymphocytes **(b)**. The datasets were: 10X_LLU, C1_FDA_HT, 10X_NCI, C1_LLU, and WaferGen_SE. The union of the top 1000 highly variable genes (HVG) of five data sets was used as the gene set for batch correction. Batch effect corrections were performed using scRNA-seq data derived from spiked-in mixed cells in which either 5% or 10% cancer cells were spiked into the sample B cells. Four data sets were analyzed: 10X_Mix10%_LLU, 10X_Mix5%_NCI, 10X_Mix5%_F1_NCI, 10X_Mix5%_F2_NCI. The top 1000 HVG were used for batch correction. Batch effect corrections were performed using four scRNA-seq data sets containing biologically distinct cells, including two breast cancer cell (A) datasets (10X_LLU_A and 10X_NCI_A) and two normal B cell (B) datasets (10X_LLU_B and 10X_NCI_B). The top 1000 HVG were used for batch correction. **(c)** Batch corrections were performed using two scRNA-seq data sets derived from samples that shared a large portion of same biological population of cells but contained a small portion of biologically distinct cells. The two data sets were: one normal cell (B) dataset (10X_LLU_B) and one spike in dataset (10X_LLU_spikein_10%A) which had 90% normal cells (B) spiked-in with 10% of cancel (A) cells. There were large portion of subpopulation of cells shared by the two data sets. **(d)** Batch corrections were performed using two scRNA-seq data sets derived from samples that contained a large portion of biologically distinct cells but shared a small portion of same biological population of cells. The two data sets were: one cancer cell (A) dataset (10X_LLU_A) and one spike_in dataset (10X_LLU_spikein_10%A) which had 90% normal cells (B) spiked-in with 10% of cancer (A) cells. The small portion of spike-in cancer cells was shared by the two datasets. Bar plots **(a-c)** showed the modified alignment score using six batch correction methods, respectively.

In all the above three scenarios (**Fig. 4a-c**), CCA out-ranked the other methods according to the modified alignment score. However, based on t-SNE and UMAP, CCA, MNN, and BBKNN all performed fairly well for the breast cancer cells (sample A) (**Fig. 4a & Suppl. Fig. 10a**), B cells (sample B) (**Fig. 4b & Suppl. Fig. 10b**), or spiked in samples (**Fig. 4c & Suppl. Fig. 10c**); CCA was slightly better than the others (**Fig. 4a-c & Suppl. Fig. 10a-c**). For the breast cancer cells, MNN and BBKNN were better than limma and ComBat, but for normal B cells (sample B), CCA and MNN perform slightly better than BBKNN; limma and ComBat seemed to perform fairly well too, based on both t-SNE and alignment score (**Fig. 4b**). Scanorama seemed not to work well for the scRNA-seq datasets derived from breast cancer cells (**Fig. 4a**) or B cells across all platforms (**Fig. 4b**). The t-SNE plots in **Figure 4c** illustrate the suitability of single-cell batch correction methods applied to a scenario when a sample consisted of mixtures of distinct cell types such as the spiked-in 10X Genomics scRNA-seq data. In the spike-in data sets, CCA, MMN, Scanorama, and BBKNN all separated the spiked in cancer cells from B cells really well, whereas limma and ComBat did not, based on t-SNE plots (**Fig. 4c**). We further calculated the modified alignment scores using different sets of HVG: the top 100, 500, 2000, and 4000 using all six batch correction methods. We observed a similar, consistent pattern of performance as shown in **Fig. 4a-c,** in which the top 1000 HVG were used, except for a lower score for MNN in sample B when the top 4000 HVG were used (**Suppl. Fig. 11a-c**). Overall, CCA, MNN and BBKNN all corrected the batch effects very well in the three scenarios.

Third, we evaluated the six batch correction methods in the following scenarios consisting of 10X Genomics datasets only: (1) two biologically distinct cell types, i.e., cancer cells (10X_LLU_A & 10X_NCI_A) plus B cells (10X_LLU_B & 10X_NCI_B) only (**Fig. 4d**); (2) two biologically distinct cell types but including a spike-in sample which shared a large portion of the same population of cells, i.e., B cells (10X_LLU_B) plus 10X_LLU_spikein_10%A (**Fig. 4e**); and (3) two biologically distinct cell types but including a spike-in sample which shares a small portion of the same cells, i.e., 10X_LLU_A plus 10X_LLU_spikein_10%A (**Fig. 4f**). Considering that BBKNN has an UMAP embedded in the pipeline, we also generated a set of UMAP figures as a comparison to t-SNE plots. In scenario # (1), both t-SNE and UMAP plots showed that CCA over-corrected the batch effect as the cancer cells were not separated from the B cells. Instead, the two cell types were totally mixed together. limma and ComBat were also unable to separate sample A from sample B. In contrast, Scanorama and BBKNN worked fairly well with both t-SNE and UMAP (**Fig. 4d & Suppl. Fig. 12a**), whereas MNN did not work with either t-SNE or UMAP. This was expected, since this method requires the batches compared to share a subpopulation of highly similar cells, which breast cancer cells and B lymphocytes do not; there was no spike-in to provide a portion of the same cell type (**Fig. 4d**). In scenario # (2), CCA again over-corrected batch effects as it was incapable of separating the spiked in cancer cells from B cells, whereas MNN, Scanorama, BBKNN, limma, and ComBat were able to separate the cancer cells from normal B cells both via t-SNE and UMAP (**Fig. 4e & Suppl. Fig. 12b**). In scenario # (3), MNN, Scanorama, and BBKNN all worked fairly well in separating cancer cells from B cells, whereas CCA, limma, and ComBat all over-corrected the batch effect; cancer cells and B cells were intermingled with both tSNE and UMAP (**Fig. 4f & Suppl. Fig. 12c**).

Two new batch correction methods, a newer version of MNN, fastMNN^6^, and Harmony^16, 17^ became available recently, so we also evaluated them. As shown in the **Suppl. Fig. 13**, there was no difference in tSNE and UMAP visualizations between regular MNN and fastMNN in four different data composition scenarios. However, fastMNN took much less computation time (**Suppl. Fig. 13**). If only the 10X Genomics scRNA-seq data were used, both Harmony and Scanorama separated cancer cells from B cells well (**Suppl. Fig. 14**), but in contrast to Harmony, Scanorama failed to separate the two types of cells if all scRNA-seq data across all platforms and sites (20 sets) were used (**Suppl. Fig. 9d and 9g**).

Overall, as illustrated, CCA failed to separate cancer cells from B cells (over-correction), whereas MNN, BBKNN, and Harmony worked well in separating cancer cells from B cells when all 20-scRNA-seq data sets across all platforms and sites were included (**Suppl. Fig. 9a-h & Suppl. Data 2**). However, one prerequisite for MNN is that the samples being corrected all share at least a small percentage of cells in common such as in our spike-in samples (**Fig. 4d-f**, **Suppl. Fig. 9c and Suppl. Data 2**).

### 5. Consistency of global and cell-type specific gene expression across platforms/sites and all scRNA-seq data

We first evaluated the global gene expression consistency across different platforms/sites by calculating a pairwise Pearson correlation (**R**) on the percentage of cells (**see Methods**) that expressed 500 abundant, 500 intermediate, and 500 scarce genes, as defined by bulk RNA-seq data (**Fig. 5a-f**). To account for variable sequencing depth across different platforms/sites, we selected one of the pipelines (zUMIs for UMI based and featureCounts for non-UMI based technology) and performed down sampling to 100K reads for each. We observed a much higher Pearson correlation when using the 500 highly-expressed genes than when using 500 intermediately- or 500 scarce genes in both cell types (**Fig. 5a-f**). We also observed higher consistency between the sites using the same platform or type of technologies (i.e., 10X, WaferGen, or C1). Even in the scarce 500 genes, we observed a reasonably good Pearson correlation between sites or within either 10X 3’ or WaferGen or C1 technologies (**Fig. 5c & Fig. 5f**). However, the consistency (Pearson correlation) within 3’ technologies (10x and C1_FDA_HT) or within full-length (C1_LLU, WaferGen_PE, WaferGen_SE) platforms was not always better than that between 3’ and full-length platforms. Nevertheless, we want to caution that there might be some biases in this analysis metric since the cell numbers were very different across platforms, i.e., only 66 or 80 single cells for the C1 full-length up to a few thousand cells for the 10x platform; since the fewer cells used, the larger the variations would exist owing purely to sampling statistics.

**Figure 5.**
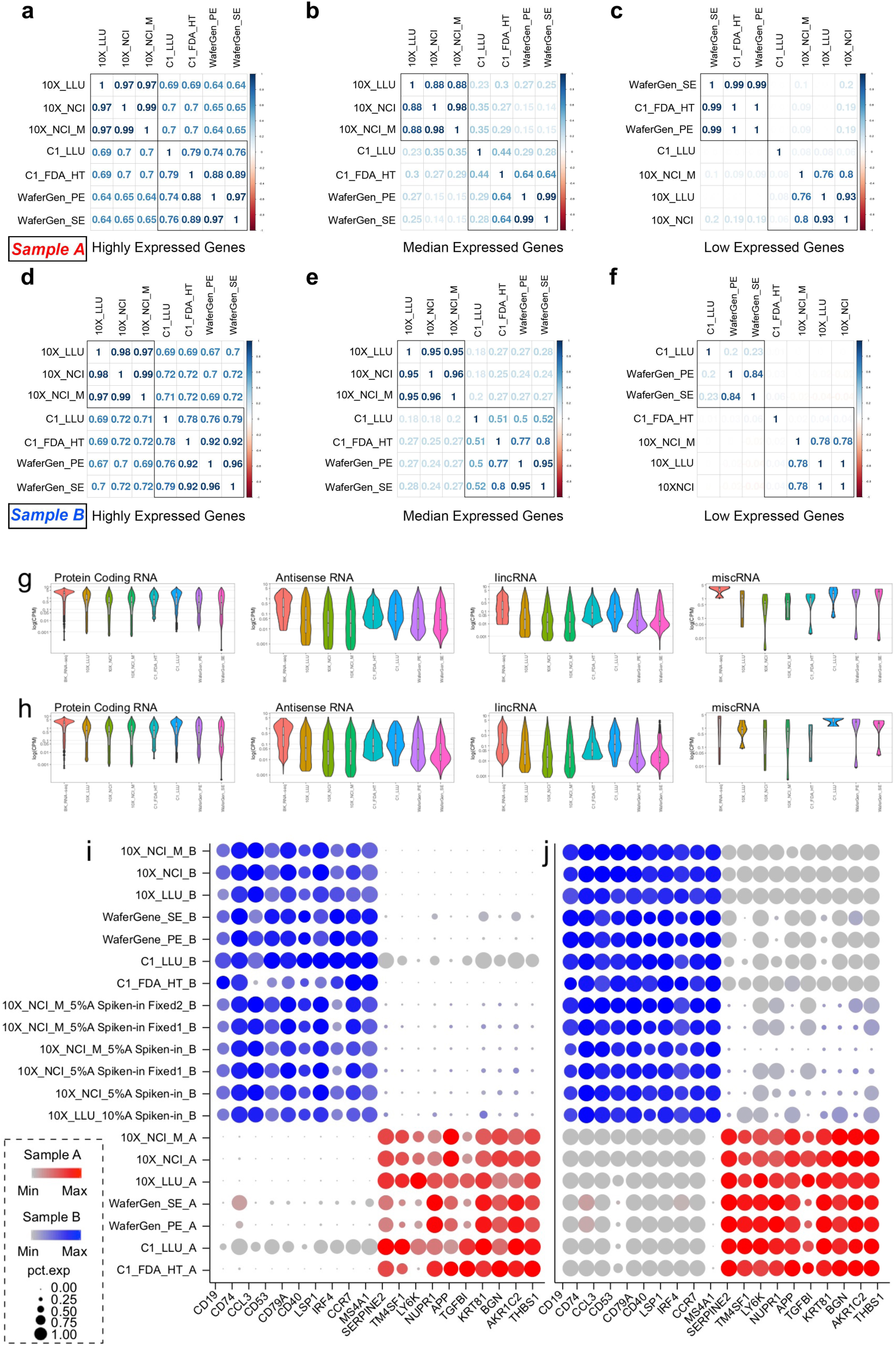
Consistency of scRNA-seq across sites/platforms and datasets. **(a-f)** Benchmarking consistency of highly abundant, intermediate and low expressed genes across all seven scRNA-seq data sets. Pairwise Pearson correlation of the percentage of cells which expressed the 500 highly-expressed genes (**a)** in cancer cells (sample A); 500 intermediately expressed genes **(b)** in cancer cells (sample A); and 500 low expressed genes **(c)** in cancer cells (sample A); the 500 most abundant genes (**d)** in B lymphocyte cells (sample B); 500 intermediately expressed genes **(e)** in B lymphocyte cells (sample B); and 500 low expressed genes **(f)** in B lymphocyte cells (sample B). **(g-h)** Benchmarking consistency of protein coding, antisense, LincRNA, and miscRNA gene expression across seven scRNA-seq datasets as well as bulk RNA-seq dataset. **(g)** Breast cancer cells (sample A) and **(h)** B lymphocytes (sample B) were profiled separately based on 4 different RNA groups across 8 different data sets. The data sets included bulk RNA-seq, 3’ scRNA-seq (10X_LLU, 10X_NCI, 10X_NCI_M, C1_FDA_HT), and full-length transcripts (C1_LLU, Waferen_PE and Wafergen_SE). **(i-j)** Split dot plots displaying ten B-cell specific vs. ten breast-cancer cell specific marker genes prior to (**i**), or post MNN batch correction (**j**). The 20 cell-type specific genes were derived from the top DEGs determined by comparing bulk RNA-seq data between breast cancer cells (sample A) and B lymphocytes (sample B). The size of each circle reflects the percentage of cells in a sample where the gene was detected, and the color indicates the average expression level within each sample. Darker color represents higher gene expression, and lighter color represents lower gene expression for the given marker gene (see inserted legends). For data sets from non-10X Genomics platform, gene expression of marker genes had a relatively low detected rate before batch effect correction. Nevertheless, after MNN batch effect correction, gene expression of marker genes was corrected well.

We then compared the single-cell gene expression profiles [log(CPM, normalized counts)] of four different RNA groups including protein coding RNA, antisense RNA, lincRNA, and miscRNA using violin plotting across all five platforms and seven datasets (**Fig. 5g-h**). As a comparison, the bulk cell RNA-seq gene expression profile was also plotted side-by-side. We noticed that WaferGen_SE gene expression profiles showed relatively higher detection sensitivity for the lower abundance transcripts. The 10X technology also seemed to show good detection (sensitivity) in the lower abundance transcripts for the protein coding RNA, antisense RNA, and lincRNA and there was high consistency across three 10X scRNA-seq datasets (10X_LLU vs. 10X_NCI vs. 10X_NCI_M). Log (CPM) gene counts across all scRNA-seq platforms and datasets for the protein coding RNA were comparable, Interestingly, using the C1 platforms (full-length and 3’), the detection range was compressed, with much lower log (CPM) values for antisense RNA and lincRNA.

We selected ten B cell-specific genes and ten breast cancer specific genes based on the ranked DEGs derived from the bulk cell RNA-seq to further evaluate consistency across all scRNA-seq platforms. **Fig. 5i** presents dot plots in which the size of each circle reflects the percentage of cells in a sample in which the gene was detected, and the color intensity reflects the average expression level within each sample. Overall, all ten B cell- and ten cancer cell-specific genes were expressed exclusively in either B cell samples or cancer cell samples except for the spiked in samples; and a lower or near noise signal detected for CD74 in B cell sample in the WaferGen_SE. We observed relatively good consistency for the B cell-specific markers^33, 34^ CD74, CD79A, LSP1, CCR7, and MS4A1 across all platforms except for C1_FDA_HT_B (**Fig. 5i**). For the cancer cell-specific markers, Serpin2, NUPR1, APP, KRT81, BGN, AKR1C2, and THBS1 were relatively consistent across platforms datasets (**Fig. 5i**). However, there were some variations and inconsistency across different platforms for some markers. We noticed relatively poor consistency or lower percentages of cells in which certain cancer cell-specific markers such as TM4SF1, LY6K, and TGFB1 were detected, particularly in WaferGen and C1_FDA_HT platforms. It is likely that a large cell-to-cell variation, i.e., endogenous biological variation, existed for these cancer cell marker genes. Particularly for platforms that capture fewer cells, sampling issues might contribute to an apparent lower percentage of cells in which these genes were detected. Interestingly, after applying MNN batch correction, we observed increased consistency in the numbers of cells and the corresponding gene expression levels detected either in B-cell or cancer cell samples for many of these genes (**Fig. 5j**).

We further exploited feature plotting and the same panel of breast cancer vs. B cell genes individually to evaluate single-cell gene expression consistency prior to and post MNN correction (**Suppl. Fig. 15a-d**). Clearly, prior to MNN batch effect correction, each of the two cell types were not clustered together and there was no clear separation between them (**Suppl. Fig. 15a & 15c**). However, after applying the MNN, cells expressing the cell-type specific marker genes were nicely clustered together and there was clear separation between two cell types (**Suppl. Fig. 15b & 15d**). Taken together, our analyses showed that choosing a batch correction algorithm appropriately to the right biological samples or scRNA-seq data sets to be analyzed is critical to improving the visualization and classification of the cell subtypes.

### 6. Single-cell detection consistency for cell-type specific markers CD40, CD74, and TPM1

We compared the consistency of single-cell gene expression across platforms for B-cell specific markers CD40, CD74, and the breast cancer cell specific marker TPM1 with a subsampling at 100k reads for each data set (**Suppl. Table 4, 5 & 6**). The B-cell specific marker gene CD40 was most often expressed at an intermediate level (1 ≤ CPM < 10) per cell, and was detected in as few as 24.9% of cells with the C1_FDA_HT to as many as 53% with the C1_LLU. A significant percentage of cells (44% to 44.6% for 10X and 23.2% to 28.1% for C1 and WaferGen) were expressed at levels close to the limit of detection (CPM < 1). In contrast, CD40 transcript was detected at low, or near noise level of (CPM <1) in breast cancer cells (**Suppl. Table 4**). However, CD74, also a B cell specific marker gene, was much more abundant (CPM ≥ 10) in almost all single cells (99.3% - 99.9%) with excellent consistency across all platforms except for C1_FDA_HT where 5% of the cells had an intermediate level (1 ≤ CPM < 10, **Suppl. Table 5**) in B cells. In contrast, CD74 was present at low or near noise levels (CPM < 1) in breast cancer cells. For this marker, full-length transcript technologies were more sensitive than the 3’-single-cell technologies (**Suppl. Table 6**). With some variation across platforms, a high percentage of single cells, expressed TPM1 in breast cancer cells, but the detection level fell mostly within an intermediate level (1 ≤ CPM < 10). In B cells, consistent with their biological nature, there was little or no detection (CPM <1) (**Suppl. Table 6**).

## Discussion

In this study, we performed a comprehensive multi-center cross-platform comparison of different scRNA-seq platforms using two well-characterized, biologically distinct reference samples. We generated 20 different scRNA-seq data sets derived from both mixed and non-mixed cell lines (**Fig. 1a and Table 1**). We applied six scRNA-seq pre-processing pipelines, seven bioinformatics normalization methods, and seven batch effect correction algorithms including CCA^7^, MNN^6^, Scanorama^8, 9^, BBKNN^10, 11^, Harmony^16, 17^, limma^35–37^, and Combat to evaluate the performance and reproducibility of scRNA-seq across 20 data sets from 4 different platforms (**Fig. 1a**). We found that there are variations in terms of number of cells and number of genes detected using different pre-processing pipelines for both UMI- and non-UMI based scRNA-seq data (**Fig. 2a & 2b**), and in addition, the gene expression (counts) correlation and the fraction of consensus genes per cell are highly variable across three UMI- and three non-UMI-based pre-processing pipelines, respectively (**Fig. 2c & 2e**). Furthermore, we benchmarked scRNA-seq performance across different platforms and testing sites using both global gene expression and some cell type specific marker genes. Using Pearson correlation based on the percentage of cells that expressed genes at different levels in each single cell, we observed a much higher consistency across different platforms and scRNA-seq datasets when using the 500 most highly-expressed genes as compared with 500 moderately abundant or 500 less abundant genes in both cancer cell and B cell samples (**Fig. 5a-f**). We also observed greater consistency between the sites within the same platform or type of technologies such as 10X, WaferGen, or C1. Even in the least abundant 500 genes for both cancer cells and B cells, we also observed a reasonably good Pearson correlation between sites or scRNA-seq data within either 10X 3’ or WaferGen and C1 technologies (**Fig. 5c & Fig. 5f**). For abundant genes such as CD74 (CPM>10), there was high consistency across platforms (∼100% of cells detected) regardless of the cell numbers captured or technologies (3’- vs. full-length). For moderately abundant transcripts (1≤ CPM <10) such as CD40 for B lymphocytes and TPM1 for breast cancer cells, there were some variations across platforms.

As mentioned previously, batch effects are a major issue when one is performing large-scale single-cell RNA-seq^6, 7^. They can result from many different sources including cell isolation and sample preparation, single cell capture, library construction, sequencing, and sampling ambiguity, etc. Most existing normalization methods were developed for bulk RNA-seq; our data showing that regressing mitochondrial genes and normalizing UMI did not remove the batch effect we observed (**Suppl. Fig. 8a/b**). This finding indicates that normalization alone cannot eliminate the batch effects when applied to scRNAseq data. Recently, several novel batch-effect-correction algorithms were developed for scRNA-seq data including CCA, MNN, Scanorama, BBKNN, and Harmony for batch-effect corrections, and k-nearest-neighbor batch-effect test (kBET) for quantification of batch effect^38^. Under certain conditions, these methods worked extremely well^8, 10^. However, our multi-platform, multicenter data sets indicated that while CCA, MNN, Scanorama, BBKNN, and Harmony worked well for batch correction of specific scRNA-seq data subsets, the nature of biological samples from which scRNA-seq data were generated from different batches, technology platforms, and sites dictated the outcome of batch effect corrections.

CCA, MNN, and BBKNN all corrected the batch variations well for the scRNA-seq data derived from biologically identical or similar samples across platforms/sites, including scRNA-seq data derived from sample A only, sample B only, or spiked-in samples only (**Fig. 4a-c**). The limma and ComBat algorithms, developed for bulk cell RNA-seq, also corrected batch effects for B cells across different platforms/sites, perhaps because these cells are more homogeneous, in which case single cell and population average data would be similar. However, for the biologically distinct samples (i.e., including both sample A and B), CCA over-corrected the batch effect and misclassified the cell types and samples, clustering both sample A cells and sample B cells together (**Fig. 4d-f, Suppl. Figs. 9b, 12a-c**). In contrast, consistent with the authors’ claims, MNN successfully removed batch effects from a variety of sources, including different platforms, laboratories, and sequencing platforms, which enabled separation of two distinct biological samples/cell types (such as sample A cells from sample B cells). However, this required a spike-in of one cell type into the other sample so that the batches being compared would have a shared subpopulation, consistent with the logic of this algorithm. This manipulation would not be necessary in situations where each of the batches being analyzed was known to contain at least one subpopulation of cells common to all the samples (**Fig. 4c, 4d-f, Supp. Fig. 9c & Suppl. Data 2)**. The poor performance of CCA when dealing with two biologically distinct samples in our analyses may be due to an insufficient amount of shared variation in gene expression between them, i.e., less heterogeneous in our two reference cell lines with somehow slightly larger heterogeneity in the breast cancer cell line cells than in B cell line cells. The primary example data set presented by the authors of CCA consisted of 13 different cell types that they identified in each batch^7^. With so many different cell types, all of which were present in each batch, the data presumably contained significant patterns of gene expression correlation that were common to all batches. These shared patterns of correlation appear to be important for the CCA algorithm. Indeed, the CCA authors stated that their Seurat procedure “uses canonical correlation analysis to identify shared correlation structures across data sets”. The shared correlation patterns, or canonical correlation vectors, are then used as a basis for scaling the data from the different batches to remove the batch effects. In our analyses, by contrast, each batch consisted of either one of the two biologically distinct cell lines or a mixture of two cell types with 90% or 95% from one or the other type. Thus, CCA might not be able to identify adequate patterns of variation in gene expression that are common to both cell lines as in our case. This hypothesis might be tested in silico by artificially generating scRNA-seq data sets with varying numbers of cell types per batch or per sample and comparing the performance of CCA with data containing different numbers of cell types.

In addition, the unique capability of UMAP^39^, an embedded bioinformatics component for BBKNN, vs. the currently most popular t-SNE visualization in separating two distinct cells types is also intriguing (**Fig. 4d, Suppl. Fig. 12a**).

In addition, our side-by-side comparison analysis comparing MNN with fastMNN did not show any differences in tSNE or UMAP visualizations using four different data sets (**Suppl. Fig. 13a-d)**. Consistent with the claim from the fastMMN authors, we found it to run very fast and use much less computation time; this is one of the major advantages of fastMNN^6^. Our analysis using Harmony^16, 17^ for batch correction showed that it works really well for scRNA-seq data regardless platforms (**Suppl. Fig. 9d & 14**), whereas Scanorama only works well for 10X scRNA-seq data, i.e., it can separate the cancer cells from B cells when all 10X scRNA-seq data across two sites including sample A, sample B, spiking-in samples were used (**Suppl. Figs. 9g & 14**). One possible explanation is that the Scanorama algorithm was originally developed using 10X Genomics scRNA-seq data^8, 16^, and it was “trained” well for 10X Genomics scRNA-seq data, but seemed to fail when other scRNA-seq platforms (such as C1 and WaferGen) were included (**Suppl. Fig. 9g)**. This result merits some further investigation.

In summary, our unique study design, which included samples of two well-characterized, biologically distinct cell lines and mixtures (either 5% or 10% of cancer cells spiked into the B cells), allowed us to benchmark the consistency and evaluate the variation of different single-cell RNA-seq technologies across different sites. We found MNN, BBKNN, and Harmony allowed correct classification of the two cell types using scRNA-seq data from all platforms and sites, regardless of single-cell technology (3’ or full-length), number of cells sequenced (thousand cells from 10X or 66 single cells from C1), or sequencing protocols (10X_NCI_M, modified shorter protocol, WaferGen SE vs. PE) (**Suppl. Figs. 9c/9d/9h and Suppl. Data 2**), suggesting the consistency of the scRNA-seq technologies as well as the high quality of the scRNA-seq data across different sites. In contrast, CCA is not able to separate two distinct biological samples, i.e., breast cancer cells from B cells. When analyzing relatively abundant genes, there was high consistency across technology platform and site, most likely due to the fact that less sampling ambiguity would be expected for these transcripts than for less abundant genes. In addition, our study provided a useful reference data set and resource for the single-cell sequencing community. We concluded that, depending on the nature of biological samples, choosing an appropriate batch effect correction algorithm and bioinformatics pipeline for scRNA-seq data analysis is critical to accurate analysis of single-cell RNA sequencing studies.

## METHODS

### Cell culture and single cell preparation

We obtained the human breast cancer cell line (HCC1395, sample A) and the matched normal B lymphocyte cell line (HCC1395 BL, sample B) from ATCC (American Type Culture Collection, VA, USA). The two cell lines were derived from the same human subject (43 years old, female). HCC1395 cells were cultured in RPMI-1640 medium supplemented with 10% FBS and 1% penicillin-streptomycin. HCC1395BL cells were cultured in IMDM medium supplemented with 20% FBS and 1% penicillin-streptomycin.

Single cell suspensions were generated by dissociating adherent cells (HCC1395) with Accutase (Innovative Cell Technologies, AT104) or by harvesting suspensions cells (HCC1395 BL). We passed all cells through a 30-micron MACS SmartStrainer (Miltenyi Biotec, 130-098-458) to filter through the cell aggregates.

### Single-cell full-length cDNA generation and RNA-seq using the C1 Fluidigm system

Single cells were loaded on a medium-sized (10-17 µm) RNA-seq integrated fluidic circuit (IFC) at a concentration of 200 cells / µl. Capture occupancy and live/dead cell at the capture site were recorded using fluorescence microscope after staining with the live/dead viability/cytotoxicity kit (Life Technologies, L3224). Full-length cDNAs were generated on the Fluidigm C1 system using the SMART-Seq v4 Ultra Low Input RNA kit (Clontech, 635026) according to the manufacturer’s protocol. Only cDNAs generated from live single cell were used for further libraries construction.

Libraries were prepared using the modified Illumina Nextera XT DNA library preparation protocol. Briefly, the concentrations of cDNAs harvested from IFC were quantified using Quant-iT PicoGreen dsDNA Assay (Life Technologies, P11496) and then further diluted into 0.1-0.3 ng / µl. 1.25 µl diluted cDNA was incubated with 1.25 µl tagmentation mix and 2.5 µl tagment DNA buffer for 10 minutes (min) at 55 °C. Tagmentation was terminated by adding 1.25 µl of NT buffer and centrifuged at 2,000 g for 5 min. Sequencing library amplification was performed using 1.25-µl Nextera XT Index primers (Illumina) and 3.75 µl Nextera PCR Master Mix in 12 PCR cycles. Barcoded libraries were purified and pooled at equal volume. Total 80 libraries were generated from HCC1395 cells (sample A) and 66 libraries were generated from HCC1395 BL cells (sample B). Library pools were sequenced on the Illumina HiSeq4000 sequencer for 150 bp paired-end sequencing.

### Single-cell 3’ End RNA-seq using C1 Fluidigm high-throughput (HT) system

High-throughput single cell 3’ end cDNA libraries were generated according to the manufacturer’s instructions. Briefly, single cells were loaded on a HT IFC at a concentration of 400 cells / µl. Capture occupancy and live/dead cell at the capture site were recorded using a fluorescence microscope after staining with live/dead viability/cytotoxicity kit (Life Technologies, L3224). After cell lysis, the captured mRNA was barcoded during the reverse transcription step with a barcoded primer, and the tagmentation step was done following the Nextera XT DNA library preparation guide. Only polyadenylated RNA containing the preamplification adapter sequence at both ends will be amplified. Lastly, sequencing adapters and Nextera indices are applied during library preparation. Only the 3’ end of the transcript was enriched following PCR amplification. 203 libraries were generated from HCC1395 cells (sample A) and 241 libraries were generated from HCC1395 BL cells (sample B). Library pools were sequenced on the Illumina NextSeq 2500, 75 bp, paired-end.

### Single-cell RNA-seq using the 10X Genomics platform

After filtering with a 30-micron MACS SmartStrainer (Miltenyi Biotec, 130-098-458), single cells were resuspended in PBS (calcium and magnesium free) containing 0.04% weight/volume BSA (400 µg/ml), and further diluted to 300 cells / µl after cell count (Countess II FL, Life Technologies). For the 5% spike-in and 10% spike-in cell mixtures, 5% or 10% of HCC1395 breast cancer cells were mixed with either 95% or 90% of HCC1395BL cells.

Single-cell RNA-seq library preparation was performed following the protocol for the 3’ scRNA-seq 10X genomics platform using v2 chemistry. Briefly, based on the cell suspension volume calculator table, 3000 cells (17.4 µl of 300 cells/ µl suspension) and barcode-beads as well as RT reagents were loaded into the Chromium Controller to generate single Gel Bead-in-Emulsions (GEMs). cDNAs were generated after GEM-RT incubation at 53 °C for 45 min and 85 °C for 5 min. cDNA amplification was performed in 12 PCR cycles following GEM cleanup. After size selection with SPRIselect Reagent, cDNA was incubated for fragmentation, end repair, A-tailing, and adapter ligation. Lastly, sequencing library amplification was performed using sample index primer in 10 cycles.

All the Libraries generated from LLU were sequenced on the NextSeq550 and HiSeq4000 with standard sequence protocol of 26×8×98 read lengths. Libraries generated from NCI were sequenced on the NextSeq550 with modified sequence protocol of 26×8×57 read lengths, and also repeated on the HiSeq4000 with standard sequence protocol of 26×8×98 read lengths.

### Single cell sequencing of fixed cells

For delayed captures, cells were fixed in methanol using a method described by Alles et al^40^. The fixed samples underwent two different treatments. For the sample 5%A Spike-in Fixed1, the normal and tumor cells were harvested, washed, counted, and a 5% spike-in mix of tumor and normal cells was prepared as described above. Approximately 130,000 cells were then processed for fixation. The cells were washed twice with 1X DPBS at 4 °C and resuspended gently in 100µl 1X DPBS (ThermoFisher Scientific, 14190144). 900ul chilled methanol (100%) was then added drop by drop to the cells with gentle vortexing. Cells were then fixed on ice for 15 mins, following which they were stored at 4 °C for 6 days. For rehydration, the fixed cells were pelleted by centrifugation at 3000 rcf for 10 mins at 4 °C and washed twice with 1X DPBS containing 1% BSA and 0.4U/µl RNase inhibitor (Sigma Aldrich, 3335399001). The cells were then counted and the concentration was adjusted to be close to 1000 cells/µl. Approximately 8000 cells were loaded for the capture onto a single cell chip for GEM generation using the 10X Genomics Chromium controller. 3’mRNA-seq gene expression libraries for Illumina sequencing were prepared using the Chromium Single Cell 3′ Library & Gel Bead Kit v2 (10X Genomics, 120237) according to the manufacturer guidelines.

For the sample 5%A Spike-in Fixed2, tumor and normal cells (approximately 4 million each) were harvested and fixed. For fixation, the cells were washed with 1X DPBS and resuspended in 10% 1X DPBS and 90% chilled methanol, as described above. Cells were then fixed on ice for 15 mins, following which they were stored at 4 °C for 24 hrs. For rehydration, the fixed cells washed with 1X DPBS containing 1% BSA and 0.4U/µl RNase inhibitor and counted. Approximately 8000 cells were loaded for the capture onto a Single cell chip for GEM generation using the 10X Genomics Chromium controller. 3’mRNA-seq gene expression libraries for Illumina sequencing were prepared using the Chromium Single Cell 3′ Library & Gel Bead Kit v2 (10X Genomics, 120237) according to the manufacturer guidelines.

### Single-cell RNA-seq using WaferGen platform

#### CELL8 Cell preparation and Single Cell Selection

A bulk cell suspension of either Cancer or BL cells (∼ 1 x 10^6^ each) was fluorescently labeled with a premade mix of Hoechst 33324 and Propidium Iodide (Ready Probes Cell Viability Imaging Kit, Thermo Fisher Scientific) in appropriate complete medium for 20 min at 37 °C. Adherent cells were first treated with Accutase as per manufacturer’s instructions (Thermo Fisher Scientific) to dissociate cells from the flask surface. Cells were washed in 1X PBS, (no Ca^2+^, Mg^2+^, Phenol Red, or serum, pH 7.4; (Thermo Fisher Scientific) centrifuged (100 X g 3 min) and resuspended in 1 mL of 1X PBS. Cell counts were determined using a Moxie Flow cell counter (ORFLO Technologies, ID, USA) and diluted to ∼1 cell in 35 nL (∼ 28,600 cells / mL) in a solution which at dispense contains: ∼0.96 of 1X PBS (1X PBS, no Ca^2+^, Mg^2+^, Phenol Red, or serum, pH 7.4; Thermo Fisher Scientific), Second Diluent (1X), RNase Inhibitor (0.4 U) and 1.92 μM of the 3’ oligo dT terminating primer: SMART-Seq® ICELL8® CDS (Takara Bio USA, CA, USA).

Each cell type solution was dispensed from a 384 well source plate into individually addressable wells in a 5,184 nanowell, 250 nL volume ICELL8 chip (SMARTer™ ICELL8® 250v Chip, Takara Bio USA, CA, USA) using a Multi Sample Nano Dispenser (MSND, SMARTer™ ICELL8® Single-Cell System, Takara Bio USA, CA, USA). Chip wells were sealed using SmartChip Optical Imaging Film (Takara Bio USA, CA, USA) and centrifuged at 300 X g for 5 min at 22 °C. All nanowells in the chip were imaged with a 4X objective using Hoechst and Texas Red excitation and emission filters. Images (TIFF format) were analyzed using automated microscopy image analysis software Cell Select (Takara Bio USA, CA, USA). The chip was stored in a chip holder at −80 °C overnight. Image analysis confirmed cell deposition followed a Poisson distribution. 600 individual nanowells, each bearing microscopy-identified single live cells, were chosen from each cell type. A well-selection map (filter file) was then autogenerated by Cell Select software to enable individual addressing of the chosen wells for addition of cDNA synthesis and library preparation reagents as detailed in the following sections. All on-chip liquid handling was performed with the MSND. After all dispensing and sealing steps, chips were centrifuged at 3,220 x g (3 min). All on-chip thermal cycling was performed using a SMARTer™ ICELL8® Thermal Cycler (Takara Bio USA, CA, USA).

#### In-chip, full-length cDNA synthesis

The ICELL8 chip (containing dispensed samples) was thawed at room temperature for 10 min and centrifuged at 3,220*g* for 3 min at 4 °C. The chip was subsequently incubated at 72 °C (3 min) and immediately placed at 4°C. Previously selected nanowells (identified as bearing a single cell via the ICELL8 filter file) were addressed with 35 nL of RT-PCR mix, and the reactions were thermally cycled in-chip as follows: (45.6 °C, 5 sec); (41 °C, 90 min); (99 °C, 9 sec); (95.5 °C, 1 min); (100 °C, 5 sec); (99 °C, 7 sec); (59 °C, 5 sec); (64 °C, 30 sec); (69.5 °C, 5 sec); (67.5 °C, 3 min); GoTo step 5 and repeat 7X, (4 °C hold).

#### In-chip, P5 index addition and tagmentation

72 primer sequences bearing P5 indices (SMART-Seq® ICELL8® Forward Indexing Primer Set A (5’-AATGATACGGCGACCACCGAGATCTACAC(*i5)*TCGTCGGCAGCGTC-3’); *i5* refers to 1-of-72 unique, 8 nucleotide indices (Hamming distance between P5 indices = 3), were dispensed from a pre-aliquoted 384-well plate in 35 nL aliquots into 72 filter-file identified, nanowell “rows”. The chip was sealed with Microseal A film and centrifuged at 3,220*g* (3 min) at 4 °C before returning to the MSND, permitting addition of Tagmentation Master Mix containing: MgCl_2_, Nextera Amplicon Tagment Mix (Illumina); Terra™ PCR Direct Polymerase Mix, and TRH (Takara Bio USA, CA, USA). The chip was sealed with Microseal A film and recentrifuged as above. Tagmentation was performed in-chip at the following temperatures: (42 °C, 4 sec); (37 °C for 30 min); (4 °C hold).

#### In-chip, P7 index and PCR reagent addition: first PCR generating 5,184 unique indices

A reagent mix containing 72 primer sequences bearing P7 indices (SMART-Seq® ICELL8® Reverse Indexing Primer Set A, 5’-CAAGCAGAAGACGGCATACGAGAT(*i7)*GTCTCGTGGG CTCGG-3’); *i7* refers to 1-of-72 unique, 8 nucleotide indices (Hamming distance between P7 indices = 3), were dispensed from the same pre-aliquoted 384-well index plate (separate location for P7 indices) in 35 nL aliquots, into 72 filter file identified “columns” of the chip. As a consequence of adding separate P5 and P7 indices to rows or columns, a 72 x 72 *m x n* matrix of combinatorial P5 and P7 pairs was generated, uniquely identifying each of the 5,184 nanowells. The chip was sealed with SmartChip Sealing Film and centrifuged at 3,220*g* for 3 minutes at 4 °C. PCR cycling was performed as follows: (77 °C, 12 sec); (72 °C, 3 min); (99 °C, 11 sec); (95.5 °C, 1 min); (100 °C, 20 sec); (99 °C, 10 sec); (53.3 °C, 5 sec); (58 °C, 15 sec); (71 °C, 5 sec); (67.5 °C, 2 min; (Go To step5 and repeat 7X); (4 °C, hold).

#### Off-chip, sample extraction and purification of round 1 PCR amplicons

Round 1 PCR amplicons were collected from the ICELL8 chip using the SMARTer ICELL8 Collection Kit: (Collection Fixture, Collection Tube and Collection Film) into a collection and storage tube as per manufacturer’s instructions (Takara Bio USA, CA, USA). 50% of the extracted library was purified twice using a 1X proportion of AMPure XP beads (Beckman Coulter) to a final volume of 14 µL in Elution Buffer, provided with the SMART-Seq ICELL8 Reagent Kit.

#### Off-chip, library amplification (2nd PCR)

Double-AMPure bead-purified, first round amplicon (14 µl, from above) was PCR amplified in a 50 µL volume of 2^nd^ PCR Mixture containing SeqAmp™ CB PCR Buffer (25 µl), 5X Primer Mix (P5 and P7 primers) and Terra™ PCR Direct Polymerase Mix 0.05 U/ µl at reaction (Takara Bio USA, CA, USA) via a thermal protocol: (98 °C, 2 min) x1; followed by 8 thermal cycles: (98 °C, 10 sec); (60 °C, 15 sec); (68 °C, 2 min). This sequencing-ready NGS library was purified using 1 round of a 1X proportion of AMPure XP beads (Beckman Coulter). The final elution volume was 17 µl in Elution Buffer.

#### NGS library MW profile

The NGS library concentration (ng / µl) was determined using a Qubit fluorometer (Thermo Fisher). Based on Qubit readings, 1 - 2 ng / µl was examined using a 2100 Bioanalyzer and a corresponding High Sensitivity DNA Kit (Agilent) to determine the MW profile of the size-selected library. The Bioanalyzer amplicon sizes ranged between 200 to 3000 bp, with an average size of 550 bp.

#### Sequencing the library

The library was diluted to 4 nM based on the above Bioanalyzer measurement and prepared for sequencing following the standard Illumina instructions for sequencing (Denature and Dilute Libraries Guide) for the Illumina NextSeq. Using a standard loading concentration, the library was sequenced on a NextSeq 550 using a high output 2×75 cycle cartridge (Illumina), including both index reads at 8 cycles each. These libraries are Nextera XT libraries with the difference being the index sequences. Libraries require dual-indices (8nt) for demultiplexing. Custom sequencing primers or PhiX are not required.

### Bulk cell RNA-seq

We isolated mRNA in bulk from HCC1395 and HCC1395 BL cells using miRNeasy Mini kit (QIAGEN, 217004), and built sequencing libraries using the NuGEN Ovation universal RNA-seq kit. Briefly, 100 ng of total RNA was reverse transcribed and then made into double stranded cDNA (ds-cDNA) by the addition of a DNA polymerase. The ds-cDNA was fragmented to ∼200 bps using the Covaris S220, and then underwent end repair to blunt the ends followed by barcoded adapter ligation. The remainder of the library preparation followed the manufacturer’s protocol. All the libraries were quantified with a TapeStation 2200 (Agilent Technologies) and Qubit 3.0 (Life Technologies). We sequenced the libraries on a NextSeq550 for 75 bp paired-end sequencing and on a HiSeq4000 for 100 bp paired-end sequencing.

## ONLINE BIOINFORMATICS METHODS

***Reference genome:*** The reference genome and transcriptome were downloaded from the 10X website as refdata-cellranger-GRCh38-1.2.0.tar.gz, which corresponds to the GRCh38 genome and Ensmebl v84 transcriptome. All the following bioinformatics data analyses are based on the above reference genome and transcriptome.

### Preprocessing of UMI based scRNA-seq data from the 10X platform

For UMI based 10X samples, three pre-processing pipelines Cell Ranger (v2.0.1), umitools ^19^(v0.5.3), and zUMIs^20^ (v0.0.5) were used to process the raw fastq data and generate gene count matrices. In the Cell Ranger pipeline, Cell Ranger count was used with all default parameter settings to generate gene count matrices. In umitools and zUMIs pipelines, reads were filtered out if phred sequence quality of cell barcode bases were < 10 or UMI bases < 10. In the zUMIs pipeline, option -d was used to perform downsampling analyses to 8 fixed depths (5k, 10k, 25k, 50k, 100k, 150k, 200k, and 250k) to generate gene count tables. With umitools, umi_tools whitelist with default parameter setting was used to generate a list of cell barcodes for downstream analysis. umi_tools extract was used to extract the cell barcodes and filter the reads (options: -- quality-filter-threshold=10 --filter-cell-barcode). STAR (v2.5.4b)^41^ was used for alignment to generate bam files containing the unique mapped reads (option: outFilterMultimapNmax 1) for gene counting. featureCounts (v1.6.1)^42^ was used to assign reads to genes and generate a BAM file (option: -R BAM). samtools (v1.3)^43^ sort and samtools index were used to generate sorted and indexed BAM files. Finally, umitools count (options: --per-gene --gene-tag=XT --per-cell -- wide-format-cell-counts) was used for the sorted BAM files to generate gene count per cell matrix.

### Preprocessing of non-UMI based scRNA-seq data from C1 and WaferGen platform

For non-UMI based samples, three pre-processing pipelines were used to process the raw fastq data and generate gene count matrices. The pipelines included trimming and filtering, alignment, and gene counting. In the trimming and filtering process, one of the three tools [Trimmomatic (v0.35)^44^, trim_galore (v0.4.1)^45^, or cutadapt (v1.9.1)^46^] was used to process the raw fastq data. Bases with quality less than 10 were trimmed from 5’ and 3’ ends of reads. Reads less than 20 bases were discarded for further analysis. STAR with default parameter settings was used for alignment to generate bam files. Three gene counting tools, featureCounts, RSEM (v1.3.0)^23^, and kallisto (v0.43.1)^22^ were used to generate gene counts per cell. All default parameter settings were used except the following: In RSEM, option --single-cell-prior was used to estimate gene expression levels for scRNA-seq data; Option of --paired-end was used if the data were paired-end fastqs; In kallisto, options -l 500 and -s 120 were used to represent estimated average fragment length and standard deviation of fragment length if the data were single-end fastqs.

### Preprocessing of bulk RNA-seq data

The preprocessing pipeline of bulk RNA-seq data included QC (FastQC v0.11.4)^47^, trimming and filtering (Trimmomatic), alignment (STAR), and gene counting (RSEM). The parameters setting in the pipeline was the same as the preprocessing pipelines used for non-UMI scRNA-seq data. In RSEM, the option --single-cell-prior was turned off for gene expression levels estimation of bulk RNA-seq data.

#### BGL and data sharing within the team

Working under the FDA single-cell sequencing consortium, to streamline fast data sharing, access, and analysis, we used the BioGenLink™ (BGL) platform from Digicon Corporation as a central repository to host the pre-processed data as described above. All data including the single-cell RNA-seq data were pre-processed at LLU and then the data were either uploaded into BGL from users’ local computers or using tools within BGL that utilized Globus, file transfer protocol (FTP), and secure copy protocol (SCP). A detailed data annotation files about all genomics data were also uploaded into the BGL. User groups and file permissions were carefully managed to prevent unauthorized access to all data. The major data analyses were carried out on each bioinformatics team member’s local computer, but there were certain analyses (see below) carried out in BGL using tools within the platform to cross-validate our bioinformatics pipelines.

### Performance of normalization methods across all datasets

We investigated some existing bulk RNA-seq normalization procedures including “Counts per Million (CPM)”, “Trimmed Mean of M values (TMM)”, “Upper Quantiles”, “DESeq” normalization implemented in the DESeq Bioconductor package and “Trimmed Mean of M values (TMM)” implemented in the edgeR. There were also methods that were specifically tailored to scRNA-seq data sets, such as scran and Linnorm etc. Both scran and Linnorm were run using default parameters.

We performed reads downsampling of each cell to two different read depths (10K and 100K per cell) for each data set and evaluated the performance of the normalization methods of two read depths per data set. Similar to the method used in the *scone* paper^49^, the metric we used to assess normalization methods was based on how well the two samples from the same cell were grouped with each other. In details, we used silhouette width, which is defined as,

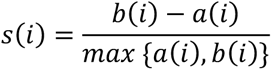

For each cell *i*, let *a(i)* be the average distance between *i* and all other cells within the same cluster. Let *b(i)* be the lowest average distance of *i* to all points in any other cluster, of which *i* is not a member. Here we defined the clustering structure that the same cells from two different sequencing runs form a single cluster, thus we have a total number of n/2 clusters if the total number of samples is n.

We calculated the silhouette width values of each dataset. The larger the silhouette width values, the better the performance of the normalization methods is.

### scRNA-seq data batch effects and batch correction pipelines

We used the gene count matrix from the Cell Ranger pipeline (10X genomics data) and STAR-featureCounts pipeline (non 10X genomics data) as input to evaluate batch correction methods. Three different conditions were considered as (1) all data sets; (2) data sets with biologically similar cells; and (3) data sets with biologically different cells. The evaluation procedure included the following four major steps:

1. Monocle2^49, 50^ strategy to filter dead cells and doublets for 10X single cell data
2. Single-cell data processing and highly variable gene (HVG) selection
3. Batch correction by six different methods
4. Evaluation by *t*-SNE or UMAP and modified alignment scores.

In step 1, all 10X single cell data sets were processed by monocle2 to filter dead cells and doublets. In monocle2, the total number of UMIs and genes for each cell were counted. The upper bound was calculated as mean plus two standard deviation (SD) and the lower bound as mean minus two SD for both the total UMIs and genes, respectively. Cells with total UMIs or genes outside of the upper and lower bounds were removed.

In the following three steps, we processed the single-cell data and selected highly variable genes for batch correction. Different data set processing and batch correction analysis strategies were developed for the three different data set scenarios and described as follows.

### Data processing and batch correction of all data sets

We examined 7 batch correction methods (CCA^7^, MNN^6^, Scanorama^8, 9^, BBKNN^10, 11^, Harmony^16, 17^, Limma^31^ and ComBat^32^) in which all 20 scRNA-seq data sets were included. After monocle2 (step 1) to remove dead cells and doublets for all 10X data sets, we used the Seurat package (v2.3.4) to process (Step 2) the 20 data sets before batch correction. Since there were large numbers of cells in the 10X data sets, we randomly sampled 1200 cells from each of the 10X data sets using the function SubsetData (Seurat package) to generate Seurat objects. For all non-10X data sets, the function CreateSeuratObject (Seurat package) was used to generate Seurat objects. The Seurat objects for all data sets were merged into one big data set. The merged data set was log transformed with the NormalizeData function and further scaled by the ScaleData function with all default parameters. Then the top HVGs (789 for CCA, 1178 for the other methods) were selected from the merged data set with the FindVariableGenes function to evaluate each batch correction method (**Suppl. Table 7**). The other procedures (Step 3 and Step 4) in the batch correction can refer to the following section on “*Data set processing and batch correction on data sets consisting of biologically similar samples and cells”*.

The following are detail descriptions of the above bioinformatics process. The gene count matrices of all data sets from the Cell Ranger pipeline (10X genomics data) and STAR-featureCounts pipeline (non-10X genomics data) were used as inputs to evaluate batch correction methods. All 20 data sets were listed as follows:

1. Sample A (7 data sets): 10X_LLU_A, 10X_NCI_A, 10X_NCI_M_A C1_FDA_HT_A, C1_LLU_A, WaferGen_SE_A and WaferGen_PE_A
2. Sample B (7 data sets): 10X_LLU_B, 10X_NCI_B, 10X_NCI_M_B, C1_FDA_HT_B, C1_LLU_B, WaferGen_SE_B, and WaferGen_PE_B
3. Spike-in sample (6 data sets): 10X_10%A_spikein_LLU, 10X_5%A_spikein_NCI, 10X_5%A_spikein_F1_NCI, 10X_5%A_spikein_NCI_M, 10X_5%A_spikein_F1_NCI_M, 10X_5%A_spikein_F2_NCI_M

The batch correction evaluation procedure included the following four major steps:

1. Filtering out low-quality cells using the Monocle2^49, 50^.
2. Single-cell data processing and highly variable gene (HVG) selection.
3. Batch correction by seven different methods.
4. Visualization using *t*-SNE, feature plot, and dot plot.

**Step 1: Filtering out low-quality cells:** For 10X Genomics data, the monocle2 strategy was first carried out to remove dead cells and doublets. In brief, the total number of UMIs and genes for each cell were counted. The upper bound was calculated as mean plus two standard deviations (SD) and the lower bound as mean minus two SD for both the total UMIs and genes, respectively. Cells with total UMIs or genes outside of the upper and lower bounds were removed.

**Step 2: Single-cell data preprocessing and highly variable gene (HVG) selection**: The Seurat package (v2.3.4) running in R version 3.5.0 were used to process the 20 data sets before the batch correction. Since there were large numbers of captured cells in the 10X data sets, we randomly down-sampled each 10X data set to 1200 cells using the Seurat function *SubsetData* and generated Seurat objects. For those non-10X data sets, the Seurat function *CreateSeuratObject* was used to generate Seurat objects. Then the Seurat objects for all the data sets were merged into one big data set. The merged data set was log transformed using the *NormalizeData* function and further scaled by *ScaleData* function with default parameters. Then the top HVGs (1178 genes for six methods except CCA) were selected from the merged data set by *FindVariableGenes* function to evaluate each batch correction method. The processed gene expression data for each sample were extracted from the Seurat data object as uncorrected data and as input for sevenbatch correction processing.

**Step 3: Batch correction using seven different methods:** In this step, seven different batch correction analysis strategies were developed as described below:

### 3.1 CCA processing of Step 2 and 3

Preprocessed data from Step 1 was used as input. Then the data sets were log transformed by the Seurat function *NormalizeData* and further scaled by the *ScaleData* function with default parameters. 500 HVGs were identified in each data set using the *FindVariableGenes* function and the union of HVGs (789 genes in total) from all data sets was used as final input data for CCA batch correction. The function *RunMultiCCA* was performed for the cross-dataset normalization and batch correction with a total of 30 estimated canonical correction vectors. The function *AlignSubspace* was used to generate the low-dimensional embedding space of each data set for visualization.

### 3.2 MNN^6^ processing of Step 3

The processed gene expression data for each sample extracted from the Seurat data object were reorganized into a data matrix, in which samples with spike-in were placed in the front of the data matrix as references. The unction *mnnCorrect* from the scran package (v1.8.4) with default parameters was carried out for running batch correction and generating MNN-corrected gene expression matrices. Then MNN-corrected data were loaded back into the Seurat data object.

### 3.3 Scanorama^8, 9^ processing of step 3

The Scanorama Python package (v0.5) was used to process the data sets and perform batch correction. The script *process.py* with the same parameters at Step1 was used to perform cell filtering and normalization. The same parameters at Step2 was then applied in the script *scanorama.py* to identify HVGs for batch correction. The function *scanorama.correct_scanpy* (from the Python package SCANPY) with default parameters was used to perform batch correction and generate Scanorama-corrected gene expression matrices.

### 3.4 BBKNN^10, 11^ processing of Step 2 and 3

The Seurat-inspired SCANPY Python workflow was applied to process the data sets. All data sets were input using the function *pd.read_csv* in the pandas package, transferred into annotated data matrices, and appended into a list using the function *anndata.AnnData* from the package anndata. Cells and genes were filtered using the functions *scanpy.api.pp.filter_cells* and *scanpy.api.pp.filter_genes* with the same parameter settings as at Step 1. The processed data matrices were merged to generate a master gene expression matrix and further log transformed and normalized by the functions *scanpy.api.pp.log1p* and *scanpy.api.pp.normalize_per_cell*. Top HVGs were identified from the merged gene expression matrix by the function *filter_genes_dispersion* with the same parameter settings at Step 2. Further log transformation (function *scanpy.api.pp.log1p*) and scaling (function *scanpy.api.pp.scale*) were performed for the newly generated gene expression matrix containing only top HVGs. The function *bbknn.bbknn* with default parameters was carried out for the batch correction.

### 3.5 Harmony processing of Step 3

Preprocessed data from Step 2 was used as input. Principal component (PC) analysis was performed by the Seurat function *RunPCA* with parameter “pcs.compute = 50”. Then Function *RunHarmony* from package “Harmony” was run for the batch correction with parameter “nclust = 50, max.iter.cluster = 100”.

### 3.6 Limma^31^ and ComBat^32^ processing of step 3

Limma and ComBat batch correction methods were applied to the uncorrected data generated at Step 2. The function *removeBatchEffect* in the limma package was carried out with default parameters for running limma batch correction and generating a limma-corrected gene expression matrix. The function *ComBat* (sva package) was run with default parameters to perform Combat batch correction and generate a ComBat-corrected gene expression matrix. Then batch corrected data were loaded back into the Seurat data object.

### Data processing and batch correction on data sets consisting of biologically similar samples and cells

We evaluated 6 batch correction methods using the samples with biologically similar cells in the following three scenarios:

1. Sample A (5 data sets, Fig. 4a): 10X_LLU_A, 10X_NCI_A, C1_FDA_HT_A, C1_LLU_A, and WaferGen_SE_A
2. Sample B (5 data sets, Fig. 4b): 10X_LLU_B, 10X_NCI_B, C1_FDA_HT_B, C1_LLU_B, and WaferGen_SE_B
3. Spike-in sample (4 data sets, Fig 4c): 10X_10%A_spikein_LLU, 10X_5%A_spikein_NCI, 10X_5%A_spikein_F1_NCI, 10X_5%A_spikein_F2_NCI

Here we used sample A as an example to describe the four major steps. **Supplementary Table 8** provides some summary information about the bioinformatics processing on sample A, sample B, and the spike-in sample.

Five data sets from sample A (9,407 cells in total) were used to evaluate batch correction methods. Monocle2 was applied to 10X_LLU and 10X_NCI data sets to remove dead cells and doublets, thus leaving a total of 8,913 cells as the input for step 2. In Step 2 and 3, we used different normalization, scaling, and gene selection methods suggested by each batch correction pipeline as described in the following.

#### CCA^15^ processing of step 2 and 3

We used Seurat (v2.3.4) to process the five data sets individually. Genes detected in fewer than 3 cells and cells containing less than 200 genes were removed from the data sets prior to further analysis. The data sets were then log transformed with the NormalizeData function and further scaled by the ScaleData function with all default parameters. The top 1,000 HVGs were identified in each data set with the FindVariableGenes function and the union of HVGs (3,203 genes in total) from the five data sets was used as final input for CCA batch correction. Depending on the number of batches, either the function RunCCA or RunMultiCCA was used to perform cross-dataset normalization and batch correction with a total of 15 estimated canonical correction vectors. The function AlignSubspace was used to generate the low-dimensional embedding space of each data set for visualization.

#### Uncorrected and MNN^6^ processing of step 2 and 3

For MNN, the scran package (v1.8.4) was used to process the data sets. Each data set was normalized through the deconvolution method^17^ by functions computeFactors and normalize with all default parameters. Gene-specific variances of each data set were calculated and decomposed into biological and technical components by the functions trendVar and decompseVar. The top 1,000 HVGs were identified by the largest 1,000 biological gene-specific variances combined across five data sets and used as input genes for cross-dataset normalization and log transformation by the functions multiBatchNorm and logcounts. The processed master gene expression matrix with the top 1,000 HVGs was considered as the uncorrected data. The function mnnCorrect with default parameters was used to perform batch correction and generate a MNN-corrected gene expression matrix.

#### fastMNN**^6^**

Each sample in the sample set was preprocessed in the same way as described for MNN. Batch effect correction was then performed using fastmnn while limiting the genes used to the highly variable gene list (subset.row=hvg.union) and setting auto.order to True. tSNE coordinates were calculated using scater (v.1.9.21) while using all available dimensions in the corrected matrix. UMAP coordinates were calculated by exporting the corrected matrix, which was imported into SCANPY (v1.3.2). Principal component analysis was performed using pp.pca with the number of principal components to compute set to the estimated number of principal components for mnnCorrect (n_comps=pcs). A neighborhood graph (pp.neighbors) was then calculated followed by calculating the UMAP coordinates (tl.umap).

#### Scanorama^8, 9^ processing of step 2 and 3

We used the Scanorama Python package (v0.5) to process the data sets and perform batch correction. The script process.py with default parameters was used to perform cell filtering and normalization. We modified the default parameter of HVG in the script scanorama.py to identify 1,000 HVGs for batch correction. Function correct (from the script scanorama.py) with default parameters was used to perform batch correction and generate a Scanorama-corrected gene expression matrix.

#### BBKNN^10, 11^ processing of step 2 and 3

We used the Seurat-inspired SCANPY Python workflow to process the data sets. The five data sets were read as annotated data matrices using the AnnData function found in the annData package and appended into a list. Cells and genes were filtered using the scanpy.api.pp.filter_cells and scanpy.api.pp.filter_genes functions with the same parameter settings as in CCA processing. The processed data matrices were merged to generate a master gene expression matrix and further log transformed and normalized by the functions scanpy.api.pp.log1p and scanpy.api.pp.normalize_per_cell. The top 1,000 HVGs were identified from the merged gene expression matrix using the function filter_genes_dispersion. Further log transformation (function scanpy.api.pp.log1p) and scaling (function scanpy.api.pp.scale) were performed on the newly generated gene expression matrix containing only the top 1,000 HVGs. The function bbknn.bbknn with default parameters was used to perform batch correction.

#### Harmony^16, 17^

We used Seurat (v2.3.4) to process the full data sets. The pre-processed Seurat data were used as inputs for Harmony batch correction analysis^16^. A sub-Seurat data object for each dataset was generated, normalized, and applied to identify the HVGs using the same method for no batch effect corrections. PCA analysis of HVGs using the function “RunPCA” was then carried out, and the first 100 PCAs were selected. The “RunHarmony” function was run for the batch effect correction. The “RunTSNE” function with parameter “reduction.use = “harmony” was finally used to plot the harmony outputs for the visualization.

#### Limma^31^ and ComBat^32^ processing of step 2 and 3

We performed limma and ComBat batch correction on the uncorrected data generated by MNN processing. The function removeBatchEffect (limma package) was used with default parameters to perform limma batch correction and generate a limma-corrected gene expression matrix. The function ComBat (sva package) was used with default parameters to perform Combat batch correction and generate a ComBat-corrected gene expression matrix.

#### *t-* SNE and UMAP plots (step 4)

The principal component (PC) analysis was performed to obtain the batch corrected gene expression matrix by either the RunPCA function (Seurat) or the calcPCA function (URD package) to estimate the number of significant PCs. We used the function Rtsne from the Rtsne package with an estimated number of PCs to generate *t*-SNE plots for uncorrected and the 5 batch corrected data except BBKNN. The function scanpy.api.tl.umap from the SCANPY python package was used with default parameters to generate UMAP plots for uncorrected and the 6 batch corrected data.

#### Modified alignment score

We adopted the idea of alignment score from Butler’s paper^7^ to calculate alignment score based on the cells’ embedding in two-dimensional space constructed by tsne or umap. However, due to the difference of cell numbers across different data sets in our study, we developed a modified alignment score calculation algorithm as follows:

1. Calculate the percentage of cells in each data set i as w_1_ (i = 1 … N, N is the total number of data sets).
2. For each cell j6j = 1 … N_j_8 of data set i, calculate how many of its k nearest-neighbors belongs to the same data set as x_ij_ and then take an average of x_ij_ in data set i to get 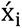 .
3. 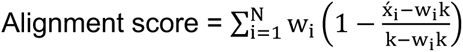
4. We chose k to be 1% of the total number of cells.

#### Data processing and batch correction on the data sets consisting of biologically distinct samples and cells

We investigated three different sample scenarios:

1. 10X_LLU_A, 10X_LLU_B, 10X_NCI_A, and 10X_NCI_B
2. 10X_LLU_A and 10X_LLU_Spikein_10%A
3. 10X_LLU_B and 10X_LLU_Spikein_10%A

**Supplementary Table 9** provides some general bioinformatics processing information for the above three different sample scenarios.

The similar four major procedures were applied to evaluate the six batch correction methods with slightly different HVG selection in uncorrected, MNN, limma, and ComBat methods. In these four methods, we identified the top 1,000 HVGs by the largest 1,000 biological gene-specific variances for each data set. Then the final HVGs were generated by taking the union of the top 1,000 HVGs from each data set for batch correction.

For Scanorama and BBKNN, we used their implemented packages on BioGenLink™ (BGL) for batch correction. The same processing and functions in Scanorama and BBKNN as described above were used to perform batch correction.

Different *t*-SNE packages were used to evaluate the six batch correction methods in biologically distinct samples and cells. Please refer to **Supplementary Table 9** for details.

#### Bioinformatics pipelines validated and performed in BGL (Biogenlink)

We carried out some bioinformatics pipelines in BGL to cross-validate some of our bioinformatics data analyses. Bioinformatics tools were created in BGL for performing batch correction of single-cell RNA-seq data using the BBKNN and Scanorama procedures and for visualizing the results of each procedure using tSNE and UMAP. For each procedure, a tool was created in BGL that allows a user to point and click to select input data and parameters for running methods from one or more packages. For each tool, BGL ran a script on the back end to execute the steps described below. Unless otherwise stated, all functions and procedures used default settings.

#### BBKNN pipeline in BGL

The BBKNN procedure ran as a Python script following the examples at weblink, https://satijalab.org/seurat/get_started.html. Data from multiple batches were read as annotated data matrices using the AnnData function found in the annData package and appended into a list. Cells and genes were filtered using the scanpy.api.pp.filter_cells and scanpy.api.pp.filter_genes functions. Data were normalized using the scanpy.api.pp.normalize_per_cell function. The scanpy.api.pp.filter_genes_dispersion function was used to identify HVGs. Data were log-transformed using the scanpy.api.pp.log1p function and scaled using the scanpy.api.pp.scale function. PCA was performed using the scanpy.api.tl.pca function. Un-corrected data were prepared for UMAP using the scanpy.api.pp.neighbors function. Data were batch-corrected and prepared for UMAP using the bbknn.bbknn function in place of the scanpy.api.pp.neighbors function. UMAP was run using the scanpy.api.tl.umap function and the results are plotted using the scanpy.api.pl.umap function.

#### Scanorama pipeline in BGL

The Scanorama procedure ran as a Python script. Data were pre-processed using the AnnData, scanpy.api.pp.filter_cells, scanpy.api.pp.filter_genes, scanpy.api.pp.normalize_per_cell, scanpy.api.pp.filter_genes_dispersion, scanpy.api.pp.log1p, and scanpy.api.pp.scale functions as described for the BBKNN procedure. Batch correction was performed using the scanorama.correct function. To prepare un-corrected and batch-corrected data for visualization, data were normalized and scaled again. PCA was performed using the scanpy.api.tl.pca function and nearest neighbors were computed using the scanpy.api.pp.neighbors function. UMAP was performed using the scanpy.api.tl.umap function and the results were plotted using the scanpy.api.pl.umap function. tSNE was performed using the scanpy.api.tl.tsne function and the results were plotted using the scanpy.api.pl.tsne function.

#### Bioinformatics evaluation of consistency of global and cell-type specific gene expression across platforms/sites and all scRNA-seq data

To investigate the consistency of global gene expression across different platforms/sites and scRNA-seq data sets, we first obtained the average gene expression (log2(TPM+1)) of bulk RNA-seq (three biological replicates) from sample A and B respectively to select benchmarking genes. We excluded the top 0.1% highly expressed genes to avoid abnormally expressed genes. We further filtered out the genes with standard deviation of gene expression greater than 1 across three replicates to obtain the robust genes. The remaining genes were used to define three different expression groups by selecting the top 500 most highly expressed, 500 intermediately expressed, and 500 infrequently expressed genes based on the ranking of average gene expression levels. For the 1500 genes selected, we calculated cell percentage per gene by defining the percentage of cells with the expressed gene (gene counts >= 1) for different scRNA-seq data sets. To get the comparable cell percentage, we only considered gene count matrices from the downsampling results (100K reads per cell) of zUMIs (10X data sets) and featureCounts (non-10X data sets) pipelines. The Pearson correlations of the cell percentage between any two scRNA-seq platforms were calculated for each of the three expression groups to evaluate the consistency.

We also examined and compared the scRNA-seq gene expression profiles across different platforms and scRNA-seq datasets based on 4 different RNA groups including protein coding RNAs, antisense RNAs, lincRNAs, and miscRNAs. The gene count matrix for each data set was used to generate the log(CPM) normalized counts. The genes which had expression of zero were removed from comparison; the filtered gene count matrices were used to extract the specific RNA group to generate violin plots.

For benchmarking marker genes across platforms, we generated dot plots of marker genes across all data sets and feature plots of individual gene by using the function SplitDotPlotGG and FeaturePlot from Seurat, respectively.

#### *t*-SNE, feature plot and dot plot

For uncorrected MNN, Scanorama, Limma and ComBat methods, the principal component (PC) analysis was first carried out using the Seurat function *RunPCA* to estimate the number of significant PCs. The Seurat function *RunTSNE* was then used to plot the significant PCs. For the CCA method, the Seurat function *RunTSNE* was applied to the 30 estimated canonical correction vectors with the parameter “reduction.use = “cca.aligned””. For the BBKNN method, the function *sc.tl.tsne* from the Python package SCANPY was applied to generate the tSNE plot. For the Harmony approach, the Seurat function *RunTSNE* was applied for the 25 harmony correction vectors with the parameter “reduction.use = “ harmony””. The function *scanpy.api.tl.umap* from the Scanpy Python package was applied with default parameters to generate UMAP plots for BBKNN and Scanorama.

For both the uncorrected and MNN-corrected data, Seurat objects from tSNE dimensional reduction were used as the data source for generating feature plots and dot plots. A total of 20 genes (10 for Sample A (Cancer cell) and 10 for Sample B (B cell)) were selected as markers for Sample A and B based on the literature. The Seurat function *SplitDotPlotGG* with default parameters was used to generate the dot plots. The Seurat function *FeaturePlot* with default parameters was run to generate the gene expression feature plots, in which each cell was colored based on the expression level of the selected gene.

#### Bioinformatics methods for single-cell detection consistency of cell-type specific markers CD40, CD74, and TPM1

To examine the consistency of three marker genes across different single cell platforms, we used the normalized gene expression data (CPM value) from the downsampling results (100K reads per cell) of zUMIs (10X data sets) and featureCounts (non-10X data sets) pipelines. The expression matrix of three marker genes per cell was generated. The expressed, infrequently expressed, intermediately expressed, and highly expressed cell percentages were defined by the percentage of cells with CPM > 0, 0 < CPM < 1, 1 ≤ CPM < 10, and CPM ≥10.

## Conflict of interests and disclaimer

All authors claimed no conflict of interests. The views presented in this article do not necessarily reflect current or future opinion or policy of the US Food and Drug Administration. Any mention of commercial products is for clarification and not intended as an endorsement.

## Authors’ contributions

CW and WX conceived and designed the study. CW managed the project and directed bioinformatics data analyses. CW drafted manuscript and annotated all the results. MMJ helped edit the manuscript. WC, CW, BT, MM, MMJ, AF, and AM performed single-cell culturing, single cell captures, scRNA-seq libraries and sequencing. XC, ZWY, YMZ, XJX, VC, YTB, BE, WX, UM, JL, JLL, and CW performed bioinformatics data analyses. WC, XC, ZWY, YMZ, YTB, XJX, VC, MM, AM, MMJ, and JJL prepared the methods for the manuscript. ZWY drew all the figures, WC and HC prepared the tables. CW, MMJ, WC, AF, WX, and YMZ revised the manuscript. All authors reviewed the manuscript. CW finalized and submitted the manuscript.

## Acknowledgements

The authors would like to thank Ms. Diana Ho of the LLU Center for Genomics for her great administrative support, particularly in coordinating the weekly Zoom conference calls and assistance in preparation of meeting minutes for the FDA SEQC-2 single-cell sequencing project. The authors would like to thank Dr. Zhong Chen at LLU and Jyoti Shetty at NCI for the technical assistance in performing sequencing; John Bettridge at NCI for the technical assistance in 10X Genomics scRNA-seq library preparation; Vyacheslav Furtak at FDA for library preparation; Wells Wu at the FDA/CBER Core Facility for Illumina sequencing. The authors also would like to thank Sangeetha Anandakrishnan of Takara Bio USA, Inc. for the technical assistance in Takara Bio ICELL8 single cell capture and library preparation. The genomic work carried out at the LLU Center for Genomics which was funded in part by the National Institutes of Health (NIH) grant S10OD019960 (CW), the Ardmore Institute of Health grant 2150141 (CW) and Dr. Charles A. Sims’ gift to LLU Center for Genomics.

## Software and code availability statement

We used many algorithms and codes which were published previously. All of our code is provided in Zenodo at the following DOI link: 10.5281/zenodo.3703082

## Data availability statement

The datasets generated during and/or analyzed during the current study are available in the SRA repository with the access code # (Sub4635070) and the data can be accessed when the paper is published. The following is the reviewer link which only contains metadata information per the SRA policy: https://dataview.ncbi.nlm.nih.gov/object/PRJNA504037?reviewer=mv5tvl7jnfaceln7lv354mchg2

## Supplementary Figures

**Supplementary Figure 1.**
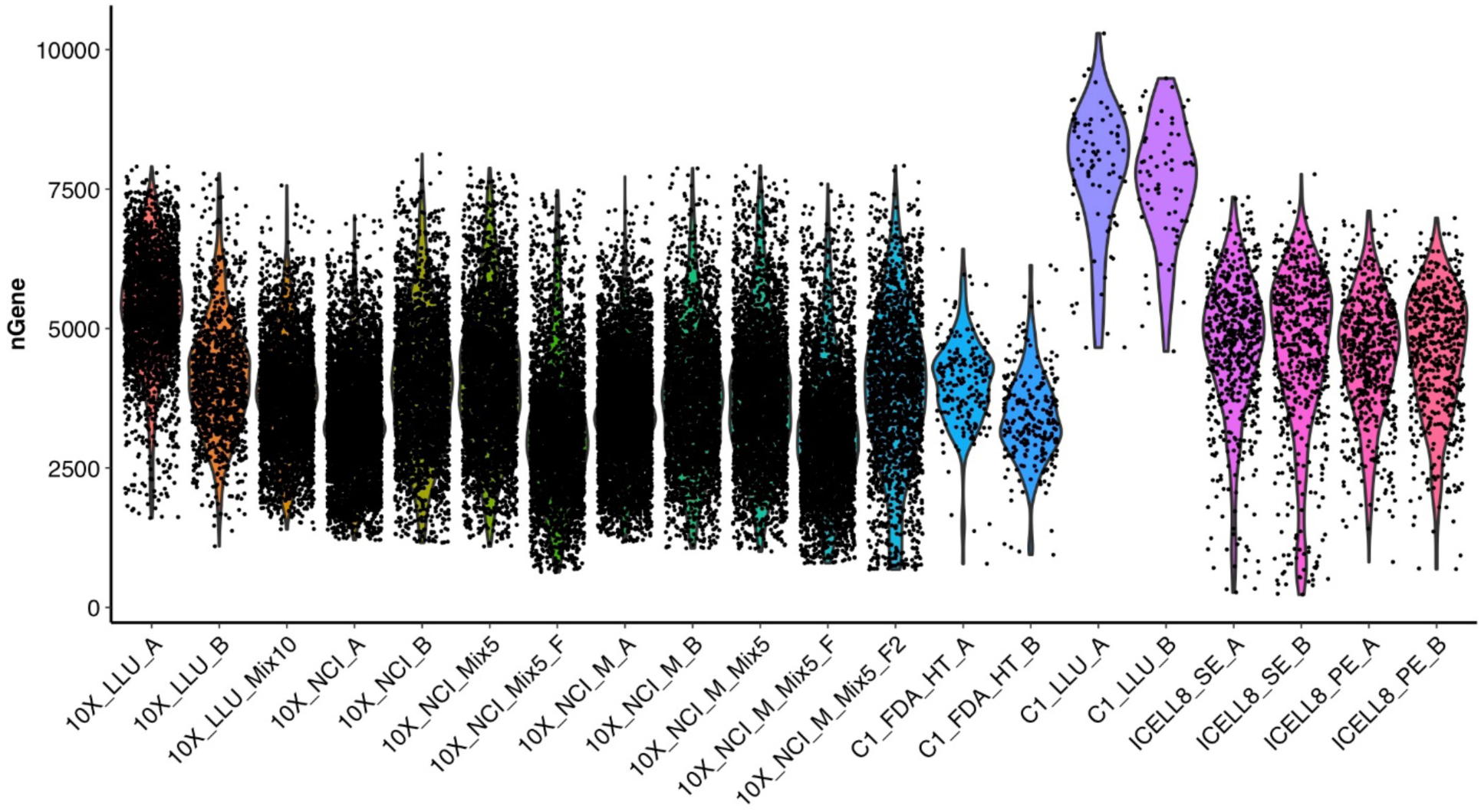
Violin plot showing the number of genes detected in each cell across all platforms/data sets. Each dot represents a cell. X-axis represents samples; Y-axis represents the number of genes detected for every cell. The shapes with color show the distributions of the data. The average number of genes detected in each cell is about 5000 and most of the cells had roughly around 2500-7500 genes, except for samples C1_LLU_A and C1_LLU_B. The 10X Genomics scRNA datasets were preprocessed using CellRanger 2.0.

**Supplementary Figure 2.**
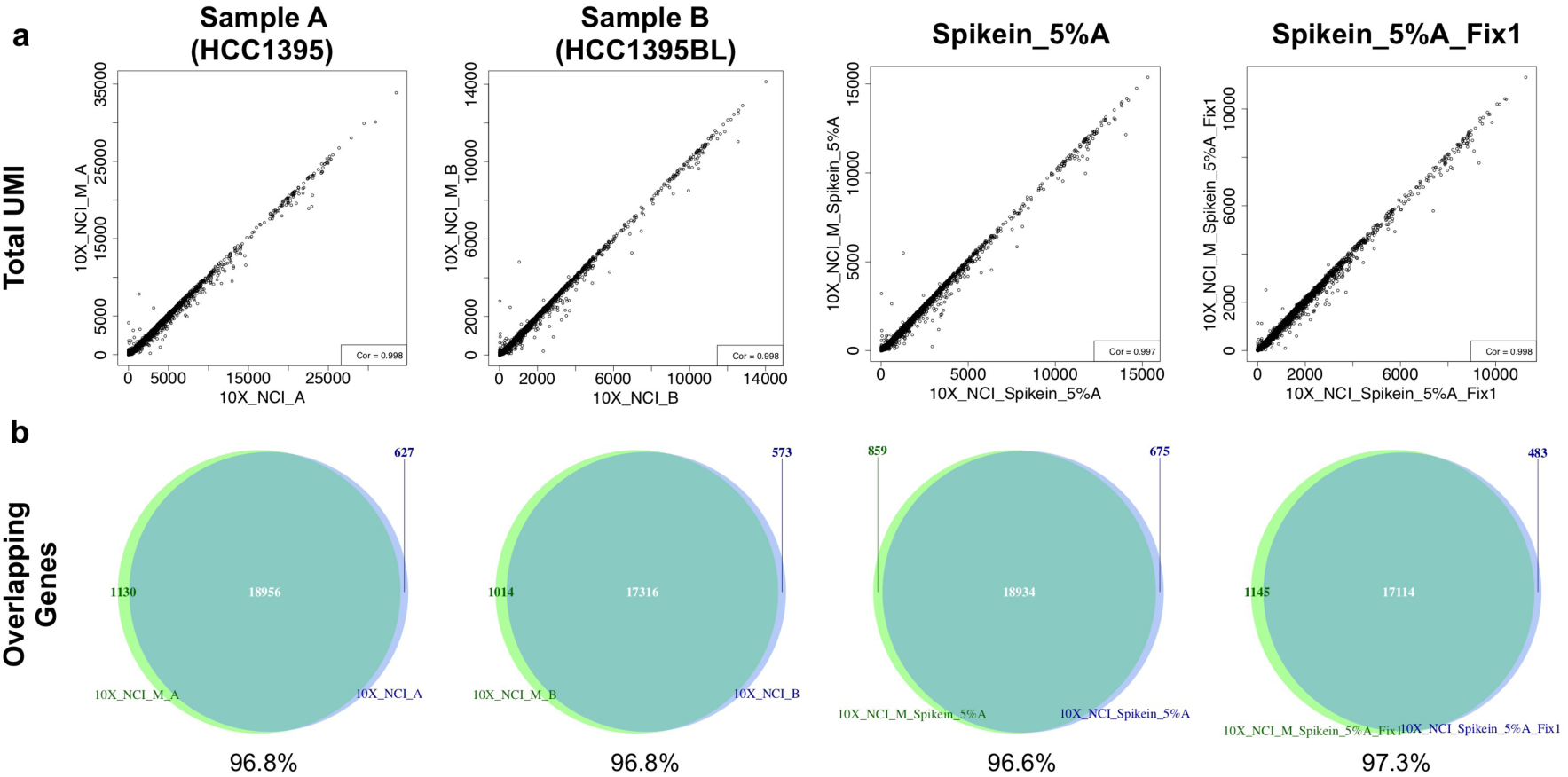
Comparison between standard and modified sequencing protocol for the 10X Genomics scRNA-seq. Correlation (**a**) between the standard and modified sequencing protocol for the 10X Genomics scRNA-seq datasets based on overlapping genes. Venn diagrams (**b)** showed the percentage of overlapping genes between two sequencing protocols. Comparisons were made between the standard (98-bp) and modified (57-bp) sequencing protocol using the overlapping genes in four different sets of libraries and eight different data sets. (**a**) The total number of unique molecular identifiers was calculated for each gene that occurred in both samples. The correlation of the data sets across all genes were then calculated for each library. (**b**) Venn diagrams showing the number of overlapping genes between two sequencing protocols. The eight data sets were: 10X_NCI_A and 10X_NCI_M_A captured from HCC1395 cancer cells, 10X_NCI_B and 10X_NCI_B_M captured from HCC1395BL normal cells, 10X_NCI_Spikein_5%A and 10X_NCI_Spikein_5%A_M captured from normal cells with 5% spike-in of cancer cells, and 10X_NCI_Spikein_5%A_Fix1 and 10X_NCI_Spikein_5%A_Fix1_M captured from normal cells with 5% spike-in of cancer cells that had been methanol fixated.

**Supplementary Figure 3.**
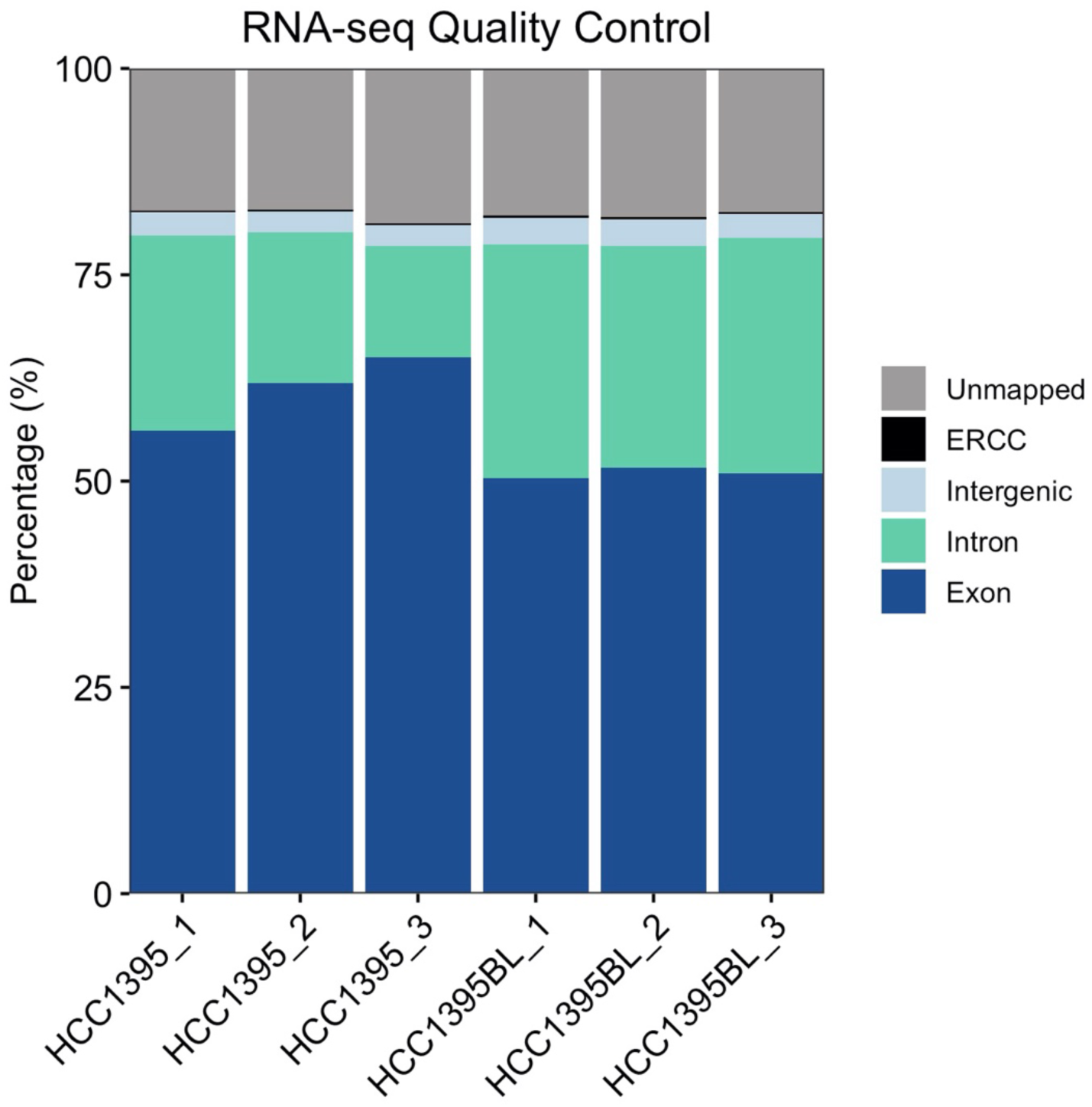
Mapping and alignment QC of bulk cell RNA-seq datasets. Bulk cell RNA-seq data sets were generated for both the HCC1395 and HCC1395BL cell lines (n=3 for each cell line). The figure shows the percentage of reads mapped to the exonic (dark blue), intronic (light green), intergenic (light blue) regions, ERCC sequences (black), or reads not mapped to the human genome (gray) in the bulk RNA-seq data.

**Supplementary Figure 4.**
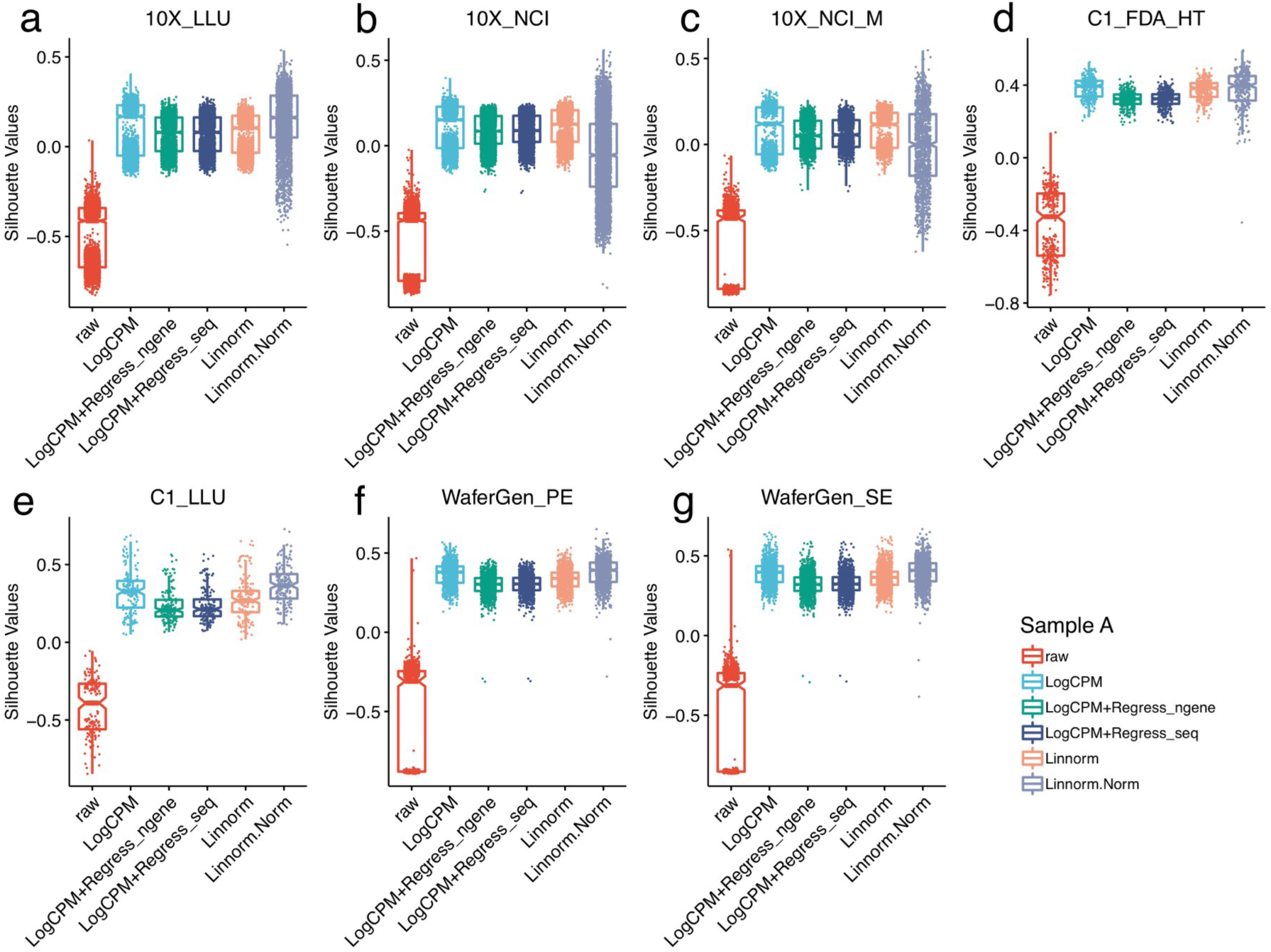
Regressing out number of detected genes does not improve the downstream silhouette scores (Sample A). (**a-g**) Boxplots of silhouette values stratified by regression-based normalization methods and 2 different Linnorm methods across 7 datasets (sample A, breast cancer cell line). Each dot represents a cell. X-axis represents normalization methods; Y-axis represents the silhouette width values for every pair of cells.

**Supplementary Figure 5.**
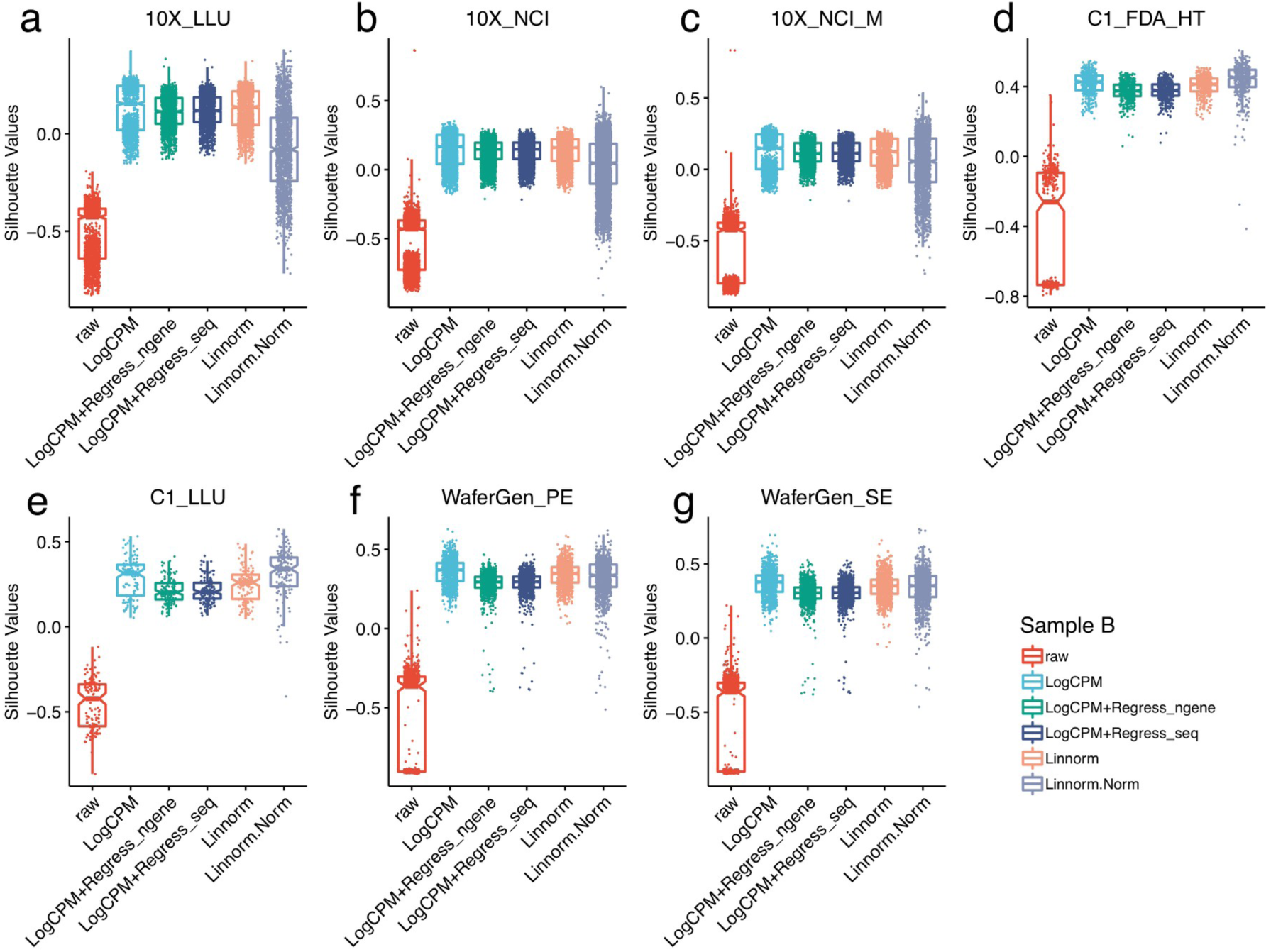
Regressing out number of detected genes does not improve the downstream silhouette scores (Sample B). (**a-g**) Boxplots of silhouette values stratified by regression-based normalization methods and 2 different Linnorm methods across 7 datasets (sample B, normal B lymphocyte cell line). Each dot represents a cell. X-axis represents normalization methods; Y-axis represents the silhouette width values for every pair of cells.

**Supplementary Figure 6.**
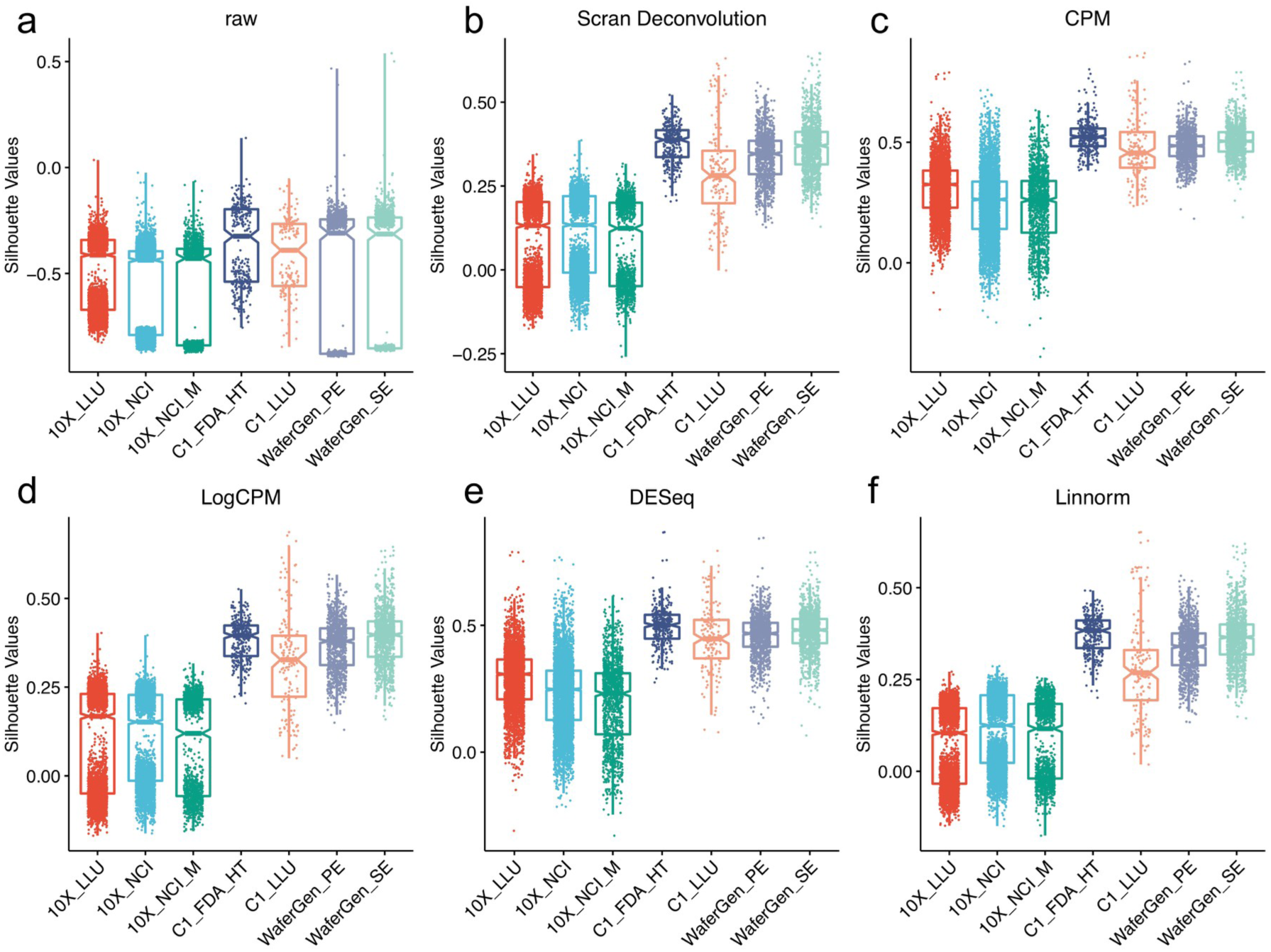
Silhouette score across different scRNA-seq platforms/data sets using different normalization methods (Sample A). **(a**) Boxplot of silhouette width values of raw gene expression across different scRNA-seq platforms/data sets; (**b-f**) Boxplots of silhouette scores across different scRNA-seq platforms/data sets using scan deconvolution, CPM, LogCPM, DESeq and Linnorm normalizations. The scores in the C1 and WaferGen platforms were consistently higher than that of the 10X data sets.

**Supplementary Figure 7.**
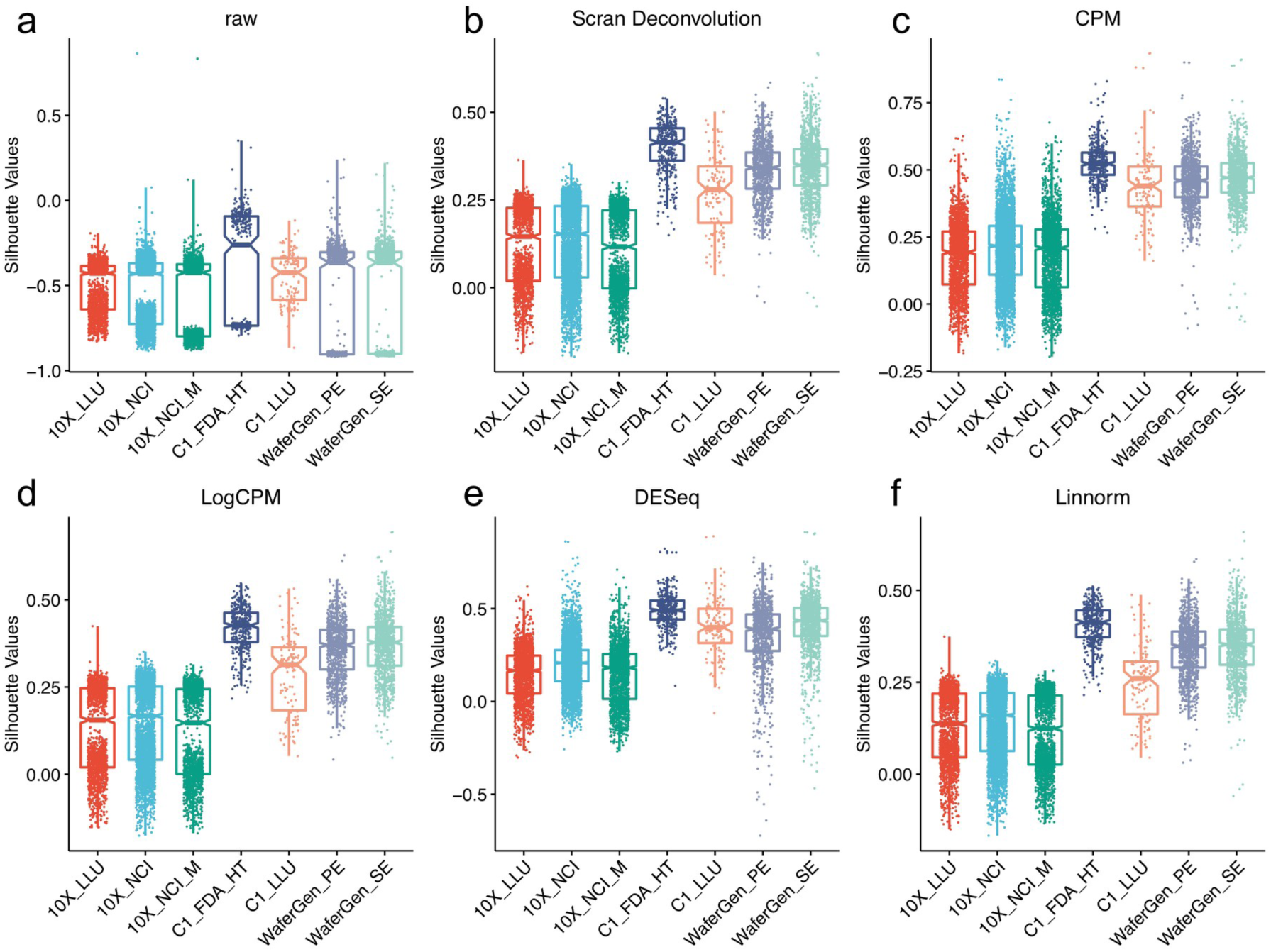
Silhouette score across different scRNA-seq platforms/data sets using different normalization methods (Sample B). (**a**) Boxplot of silhouette width values of raw gene expression across different scRNA-seq platforms/data sets (**b-f**); Boxplots of silhouette scores across different scRNA-seq platforms/data sets using scan deconvolution, CPM, LogCPM, DESeq and Linnorm normalizations. The scores in the C1 and WaferGen platforms were consistently higher than that of the10X data sets.

**Supplementary Figure 8.**
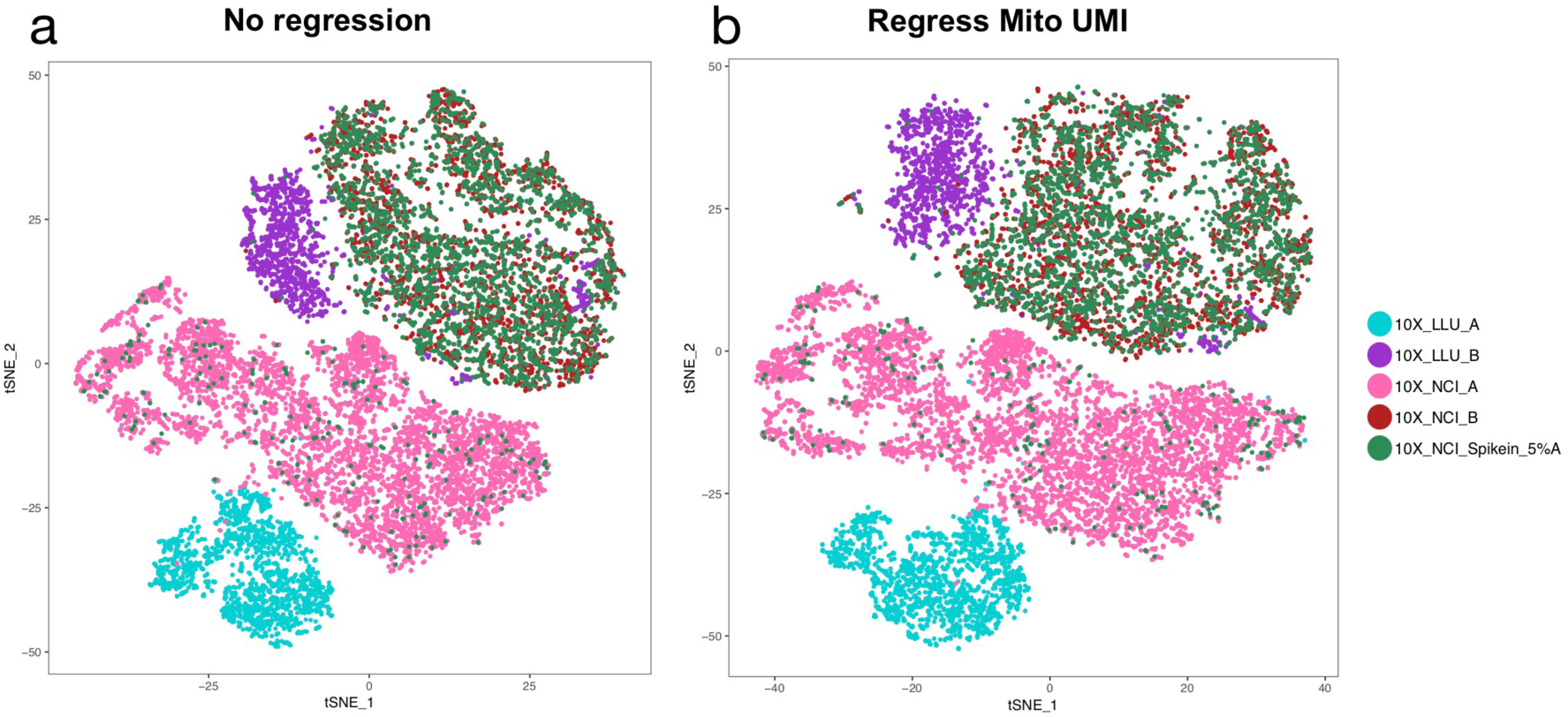
Regressing mitochondrial genes & normalizing UMI does not remove batch effects. t-SNE plots were generated from five samples sequenced at two sites after regressing out the effects of mitochondrial genes and UMI by Seurat. (a) t-SNE plot of 5 libraries/scRNA-seq data sets without mitochondrial gene regression and UMI normalization. (b) t-SNE plot of 5 libraries/scRNA-seq data sets after regression mitochondrial genes (mito) and filtering cells with mito >5%. In addition, the dataset was normalized with logNormalize. The batch effect is displayed in t-SNE plots showing that the libraries/scRNA-seq data derived from the same cell line were not clustered together. Regression mitochondrial genes and normalize dataset did not remove the observed batch effect. The five data sets are: 10X_LLU_A and 10X_NCI_A were two libraries captured from HCC1395 cancer cells; 10X_LLU_B and 10X_NCI_B were two libraries captured from HCC1395BL normal cells, and 10X_NCI_spikein_5%A was the library that had spike-in 5% of cancer cells into normal cells.

**Supplementary Figure 9.**
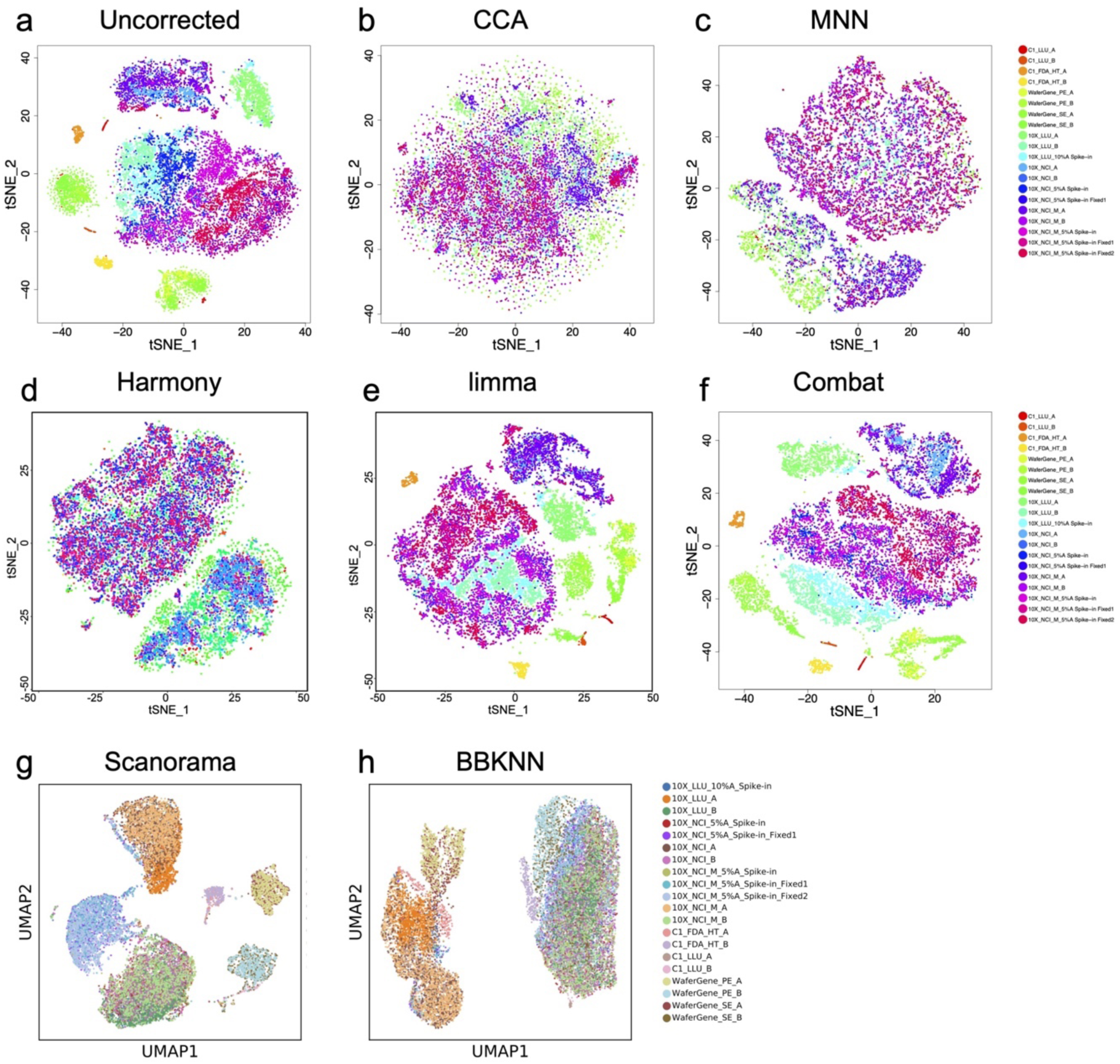
Batch-effect corrections across 20 scRNA-seq datasets. tSNE plot of 20 data sets, prior to **(a)** and post CCA (**b)**, MNN **(c)**, Harmony **(d)**, limma **(e)** and ComBat (**g),** and UMAP plot of Scanorame **(g)** and BBKNN **(h)** batch effect correction. Samples from 10X Genomics were sub-sampled to 1200 cells. Sample sharing a small portion of same biological population of cells was used as common reference. tSNE plots of 20 data sets, uncorrected (**a**), CCA (**b**), MNN (**c**), Harmony (**d**), limma (**e**) and ComBat (**f**); and UMAP plots of Scanorame (**g**) and BBKNN (**h**) batch effect correction are presented. Data set is labeled by color. MNN, Harmony and BBKNN corrected the batch variations well, in which cells were clustered into two groups as expected. After Limma, ComBat, or Scanorama batch effect correction, cells within the same cell types from different platforms or sites failed to group together. After CCA batch effect correction, cells were mixed together and not separated. Most of the methods were carried out with the R platform except for BBKNN and Scanorama, which used Python scripts.

**Supplementary Figure 10.**
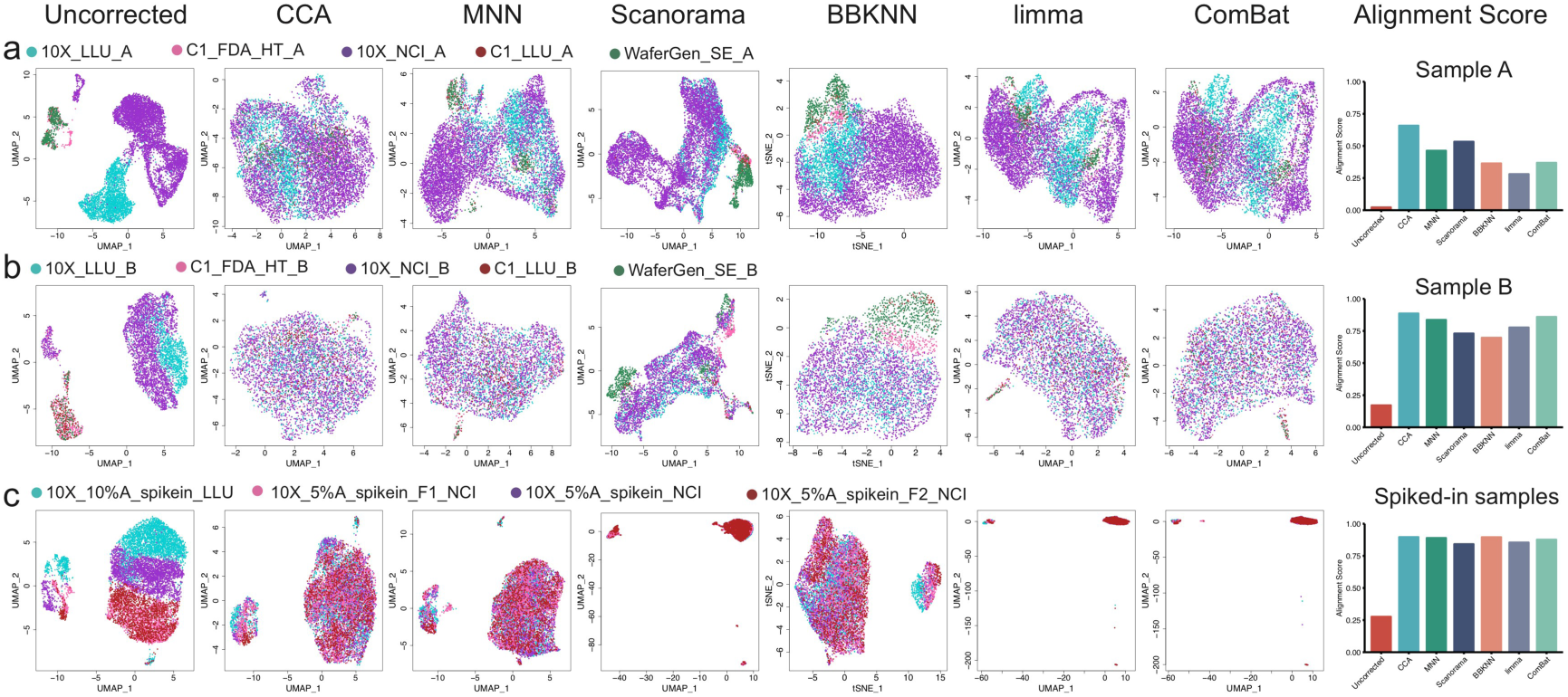
UMAP and modified alignment scores across five different data sets using six batch correction methods (top 1000 HVG). **(a)** Sample A-cancer cells; (**b**) Sample B-B-cells; **(c)** 5%A & 10%A spike-in in B-cells. Batch correction was performed using six methods on (**a**) breast cancer cells (sample A), (**b**) normal B lymphocyte cells (sample B), and (**c**) spiked-in samples where either 5% or 10% of cancer cells were spiked in into the sample B cells. The five data sets of **(a)** and (**b**) include 10X_LLU, C1_FDA_HT, 10X_NCI, C1_LLU, and WaferGen_SE. The four data sets of (**c**) include 10X_Mix10_LLU, 10X_Mix5_NCI, 10X_Mix5_F_NCI, 10X_Mix5_F2_NCI.

**Supplementary Figure 11.**
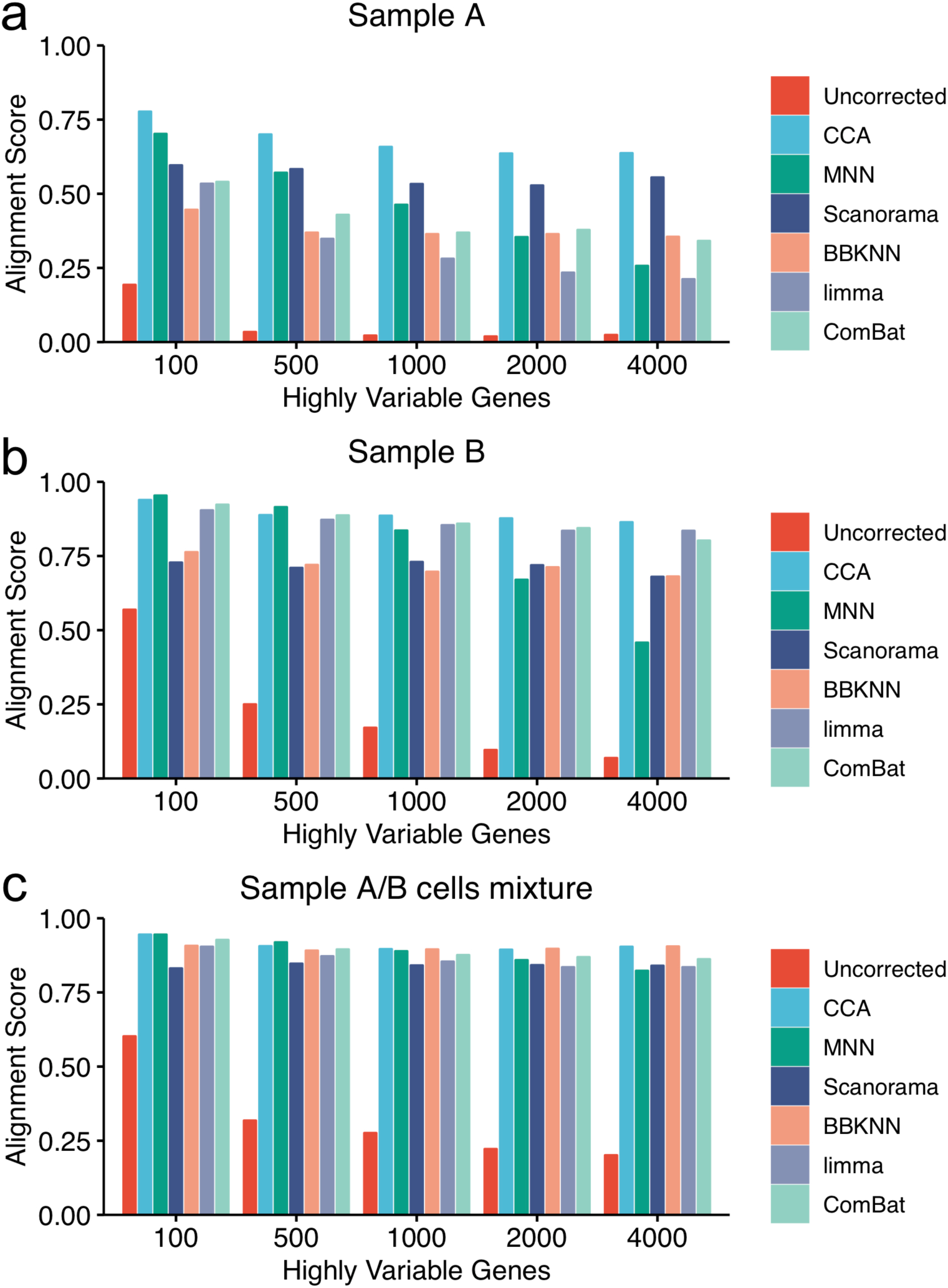
Modified alignment scores using six batch correction methods and different highly variable genes across 5 data sets. Modified alignment scores were calculated in (**a**) breast cancer cells (sample A), (**b**) normal B lymphocyte cells (sample B), and (**c**) spiked-in samples where either 5% or 10% of cancer cells were spiked in into the sample B cells. The five data sets of **(a)** and **(b)** include 10X_LLU, C1_FDA_HT, 10X_NCI, C1_LLU, and WaferGen_SE. The four data sets of (**c**) include 10X_Mix10_LLU, 10X_Mix5_NCI, 10X_Mix5_F_NCI, 10X_Mix5_F2_NCI.

**Supplementary Figure 12.**
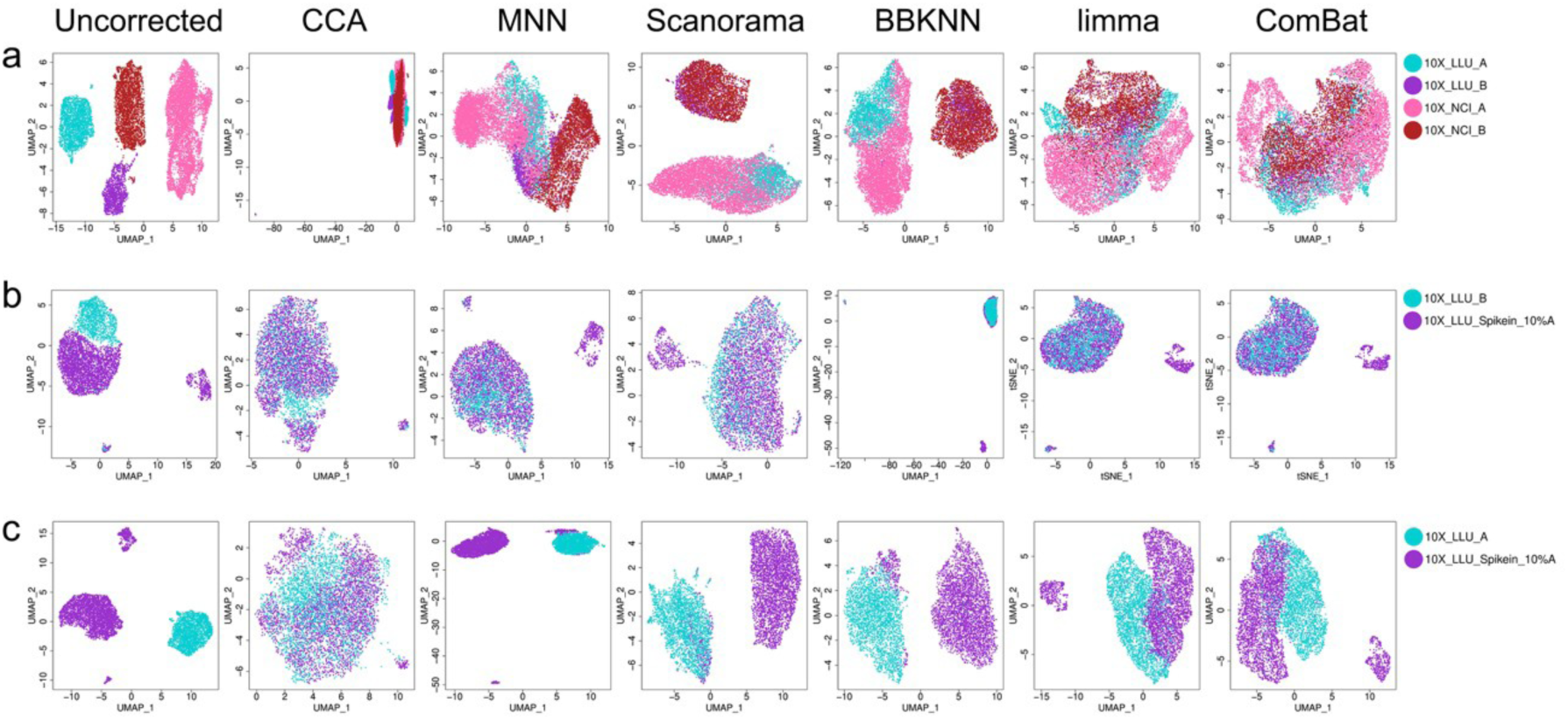
Batch effect correction visualization using UMAP plots. **(a**) Batch effect corrections were performed using four scRNA-seq data sets containing biologically distinct cells, including two breast cancer cell datasets (10X_LLU_A and 10X_NCI_A) and two normal B cell datasets (10X_LLU_B and 10X_NCI_B). (**b**) Batch corrections were performed using two scRNA-seq data sets derived from samples that shared a large portion of the same biological population of cells but contained a small portion of biologically distinct cells. The two data sets were: one normal cell dataset (10X_LLU_B) and one spike in dataset (10X_LLU_spikein_10%A). (**c**) Batch corrections were performed using two scRNA-seq data sets derived from samples that contained a large portion of biologically distinct cells but shared a small portion of the same biological population of cells. The two data sets were: one cancer cell dataset (10X_LLU_A) and one spike_in dataset (10X_LLU_spikein_10%A).

**Supplementary Figure 13a-c.**
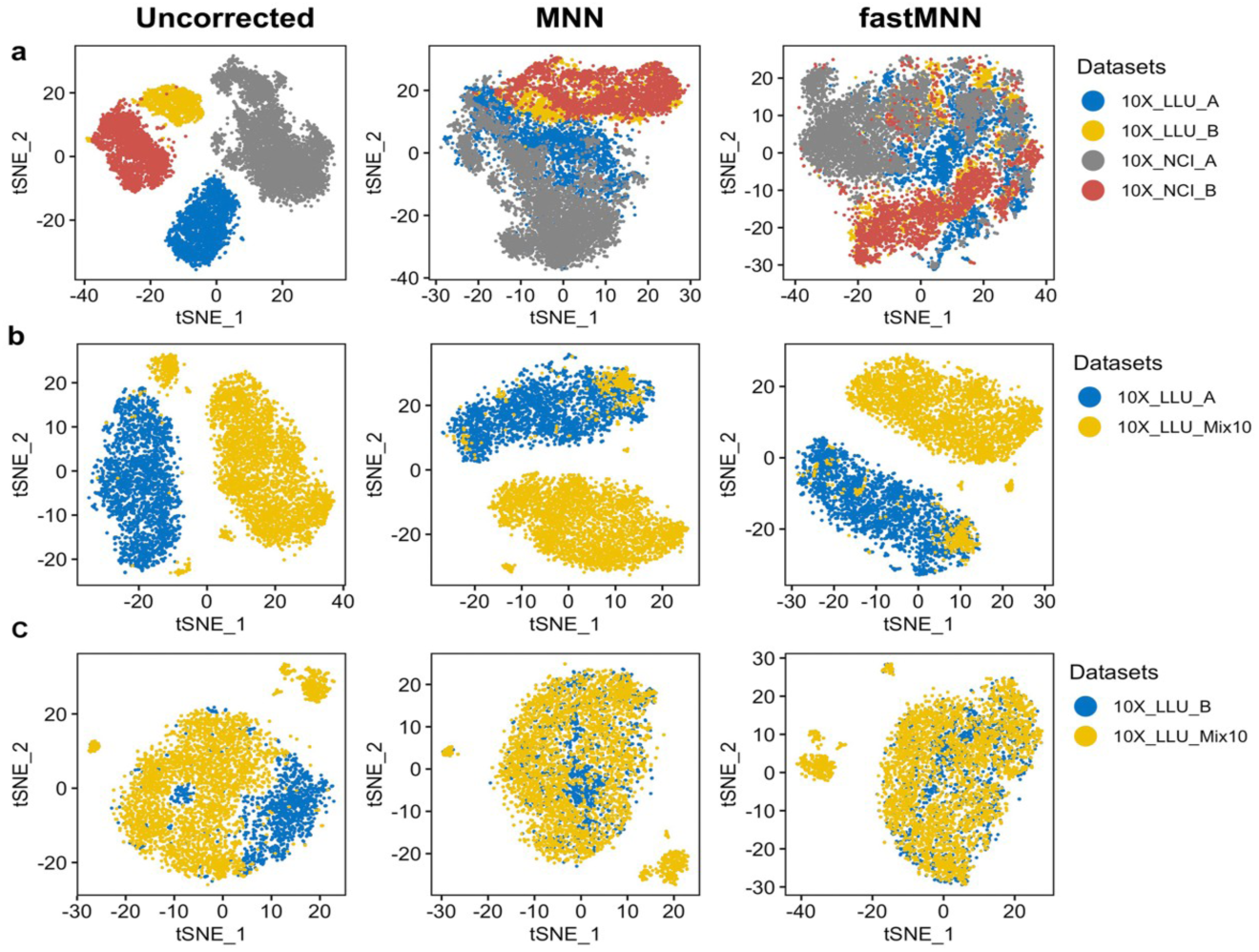
t-SNE plots of the data containing biologically distinct cells after MNN and fastMNN batch correction. (**a)** Batch effect corrections were performed using four scRNA-seq data sets containing biologically distinct cells, including two breast cancer cell datasets (10X_LLU_A and 10X_NCI_A) and two normal B cell datasets (10X_LLU_B and 10X_NCI_B). (**b**) Batch corrections were performed using two scRNA-seq data sets derived from samples that contained a large portion of biologically distinct cells but shared a small portion of same biological population of cells. The two data sets were: one cancer cell dataset (10X_LLU_A) and one spike_in dataset (10X_LLU_spikein_10%A). (**c**) Batch corrections were performed using two scRNA-seq data sets derived from samples that shared a large portion of same biological population of cells but contained a small portion of biologically distinct cells. The two data sets were: one normal cell dataset (10X_LLU_B) and one spike in dataset (10X_LLU_spikein_10%A).

**Supplementary Figure 13d-f.**
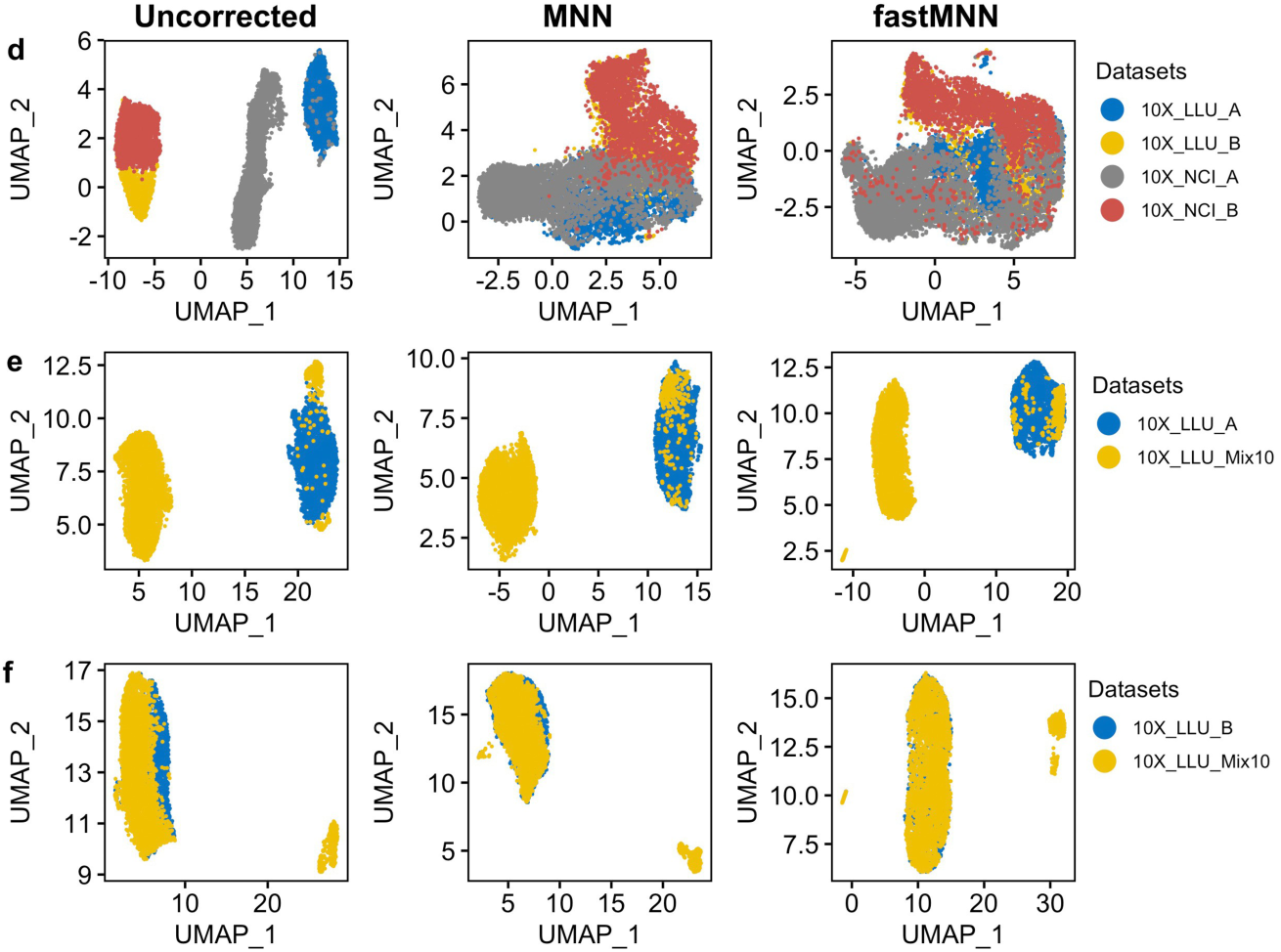
UMAP plots of the data containing biologically distinct cells after MNN and fastMNN batch correction. For UMAP plotting after batch correction, SingleCellExperiment objects were converted to Seurat object and RunUMAP/DimPlot from Seurat package with default parameters were used. (**d**) Batch effect corrections were performed using four scRNA-seq data sets containing biologically distinct cells, including two breast cancer cell datasets (10X_LLU_A and 10X_NCI_A) and two normal B cell datasets (10X_LLU_B and 10X_NCI_B). (**e**) Batch corrections were performed using two scRNA-seq data sets derived from samples that contained a large portion of biologically distinct cells but shared a small portion of same biological population of cells. The two data sets were: one cancer cell dataset (10X_LLU_A) and one spike_in dataset (10X_LLU_spikein_10%A). (**f**) Batch corrections were performed using two scRNA-seq data sets derived from samples that shared a large portion of same biological population of cells but contained a small portion of biologically distinct cells. The two data sets were: one normal cell dataset (10X_LLU_B) and one spike in dataset (10X_LLU_spikein_10%A).

**Supplementary Figure 14.**
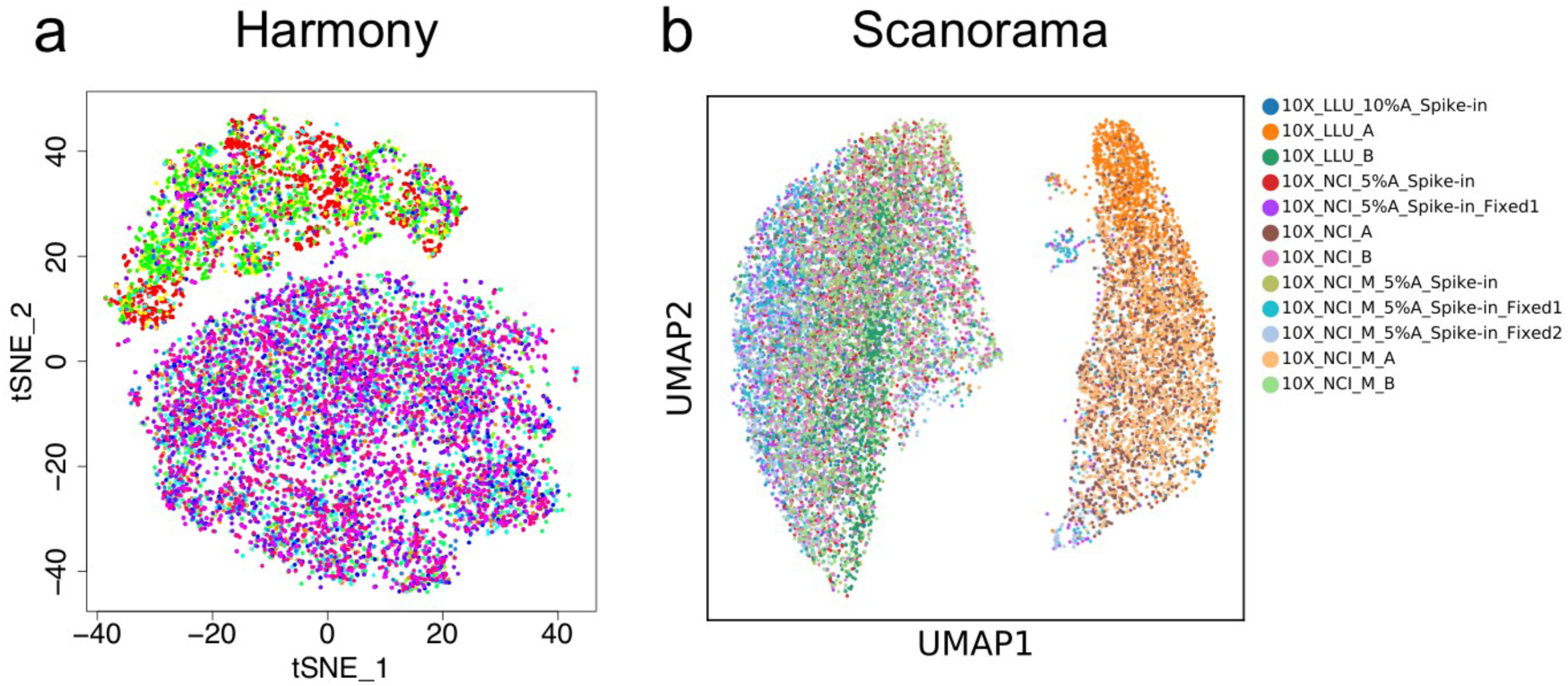
Harmony and Scanorama batch effect corrections using 10X Genomics scRNA-seq datasets from two centers. (a) t-SNE plot of 12 data sets post Harmony batch correction. (**b**) UMAP plot of 12 data sets post Scanorama batch correction. All the data sets were processed with subsampling to 1200 cells, using one common reference.

**Supplementary Figure 15a-b.**
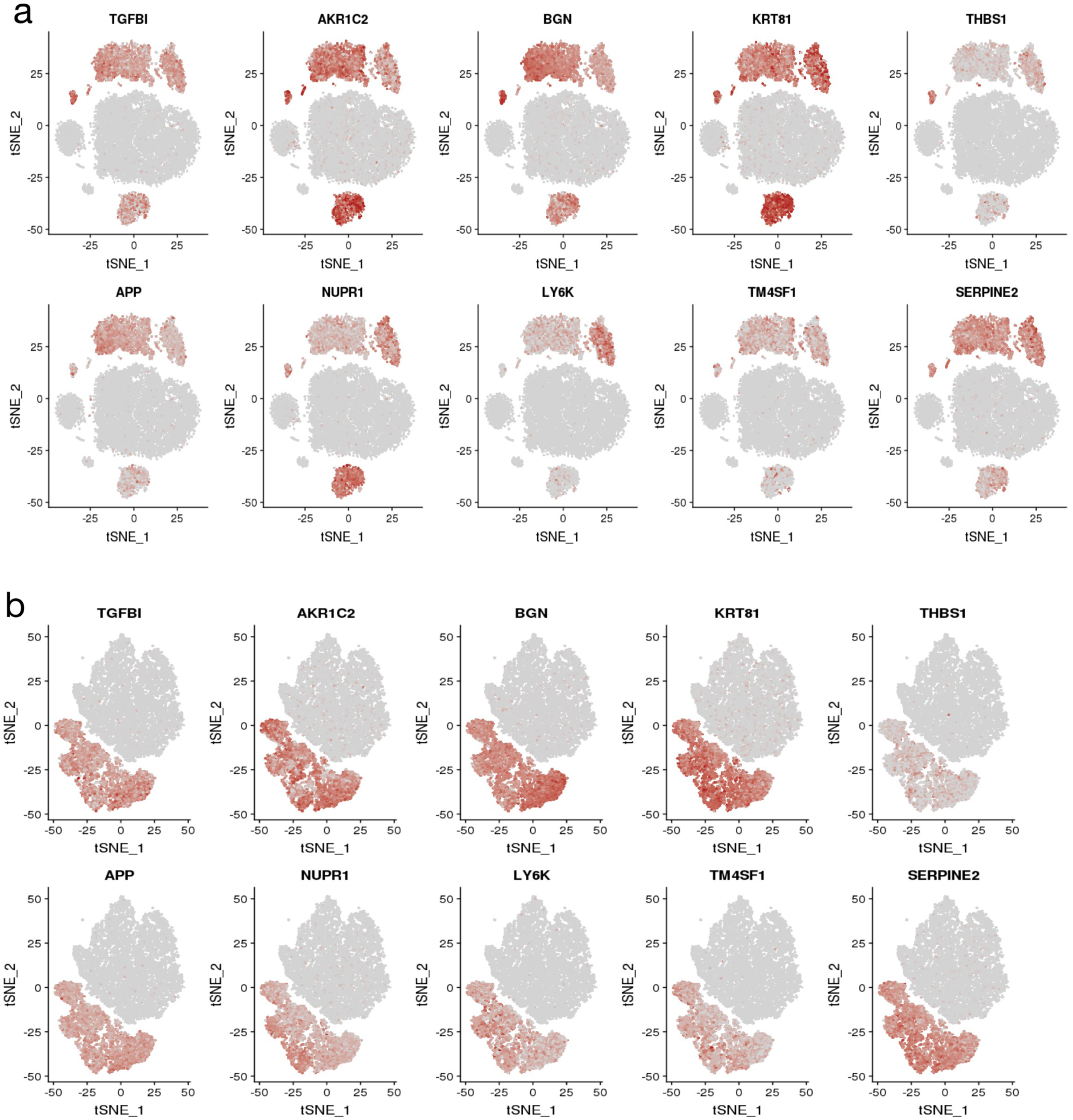
Cell type-specific marker genes for breast cancer cells (Sample A) across all scRNA-seq platforms/data sets. Feature plots of top 10 up/down- regulated genes for 20 data sets pre (**a**) and post (b) MNN batch effect correction. Genes with relatively high expression levels in each cell are highlighted in brick red in tSNE plot, which allows the visualization of the expression of a particular gene in the context of all the cells examined and also helps validate the specificity of the marker or the quality of the clustering. Before batch correction (**a**), cells expressing marker genes didn’t group together; but after MNN correction (**b**), cells expressing marker genes grouped together very well.

**Supplementary Figure 15c-d.**
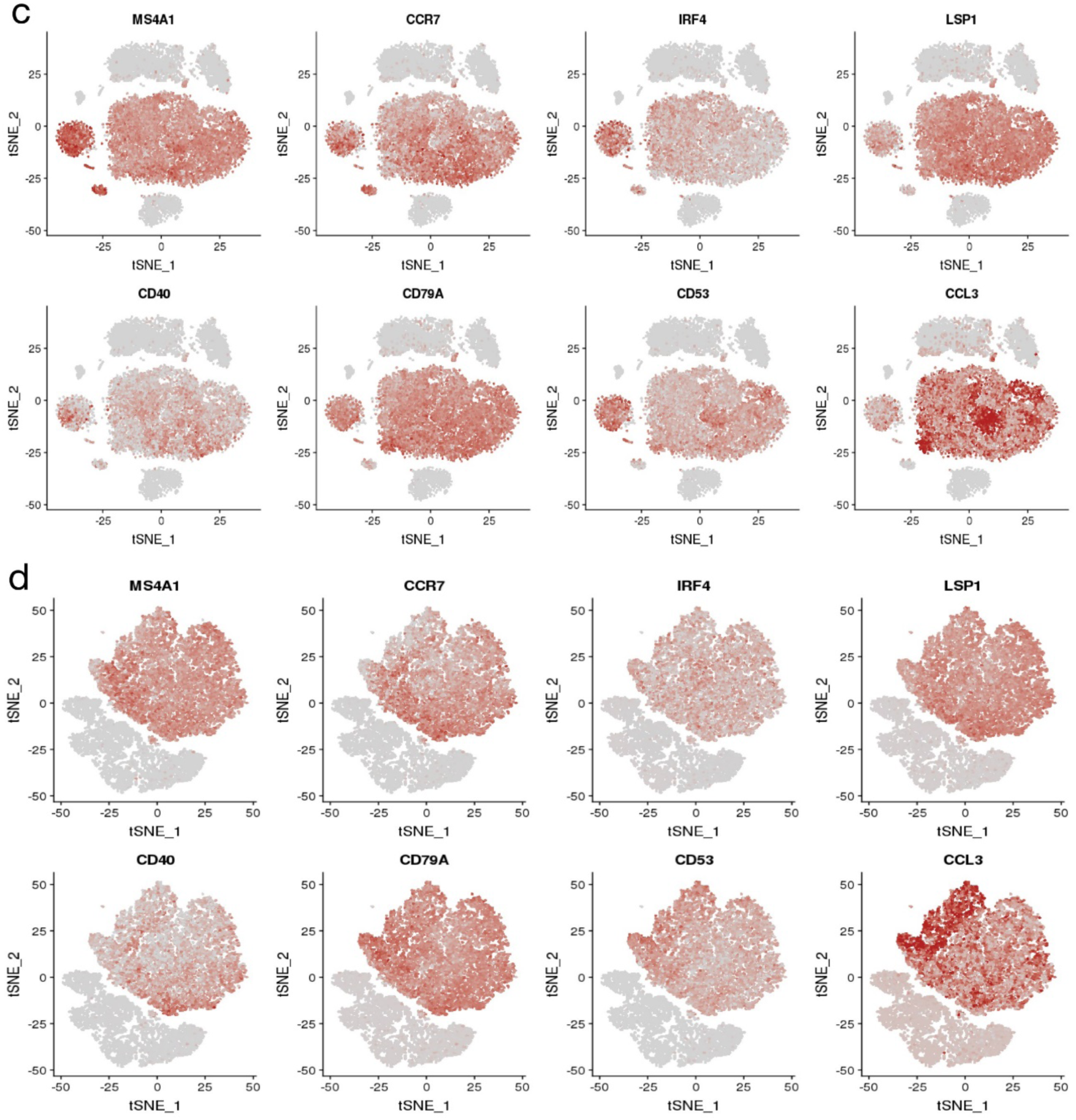
Cell type-specific marker genes for normal B lymphocytes (Sample B) across all scRNA-seq platforms/data sets. Feature plots of top 10 up/down-regulated genes for 20 data sets pre (**c**) and post (**d**) MNN batch effect correction. Genes with relatively high expression levels in each cell are highlighted in brick red in tSNE plot, which allows the visualization of the expression of a particular gene in the context of all the cells examined and also helps validate the specificity of the marker or the quality of the clustering. Before batch correction (**c**), cells expressing marker genes were not clustered together; but after MNN correction (**d**), cells expressing marker genes grouped together very well.

